# Metatranscriptomics-based metabolic modeling of patient-specific urinary microbiome during infection

**DOI:** 10.1101/2024.03.25.586446

**Authors:** Jonathan Josephs-Spaulding, Hannah Clara Rettig, Johannes Zimmermann, Mariam Chkonia, Alexander Mischnik, Sören Franzenburg, Simon Graspeuntner, Jan Rupp, Christoph Kaleta

## Abstract

Urinary tract infections (UTIs) are a major health concern which incur significant socioeconomic costs in addition to substantial antibiotic prescriptions, thereby accelerating the emergence of antibiotic resistance. To address the challenge of antibiotic-resistant UTIs, our systems biology approach uncovers patient-specific uromicrobiome insights that are focused on community utilization of metabolites. By leveraging the distinct metabolic traits of patient-specific pathogens, we aim to identify metabolic dependencies of pathogens and provide suggestions for targeted interventions for future studies. Combining patient-specific metatranscriptomic data with genome-scale metabolic modeling and data from the Human Urine Metabolome, this study explores UTIs from a systems biology perspective through the reconstruction of tailored microbial community models to mirror the metabolic profiles of individual UTI patients’ urinary microbiomes. Delving into patient-specific bacterial gene expressions and microbial interactions, we identify metabolic signatures and propose mechanisms for UTI pathology. Our research underscores the potential of integrating metatranscriptomic data using systems biological approaches, providing insights into disease metabolic mechanisms and potential phenotypic manifestations. This contribution introduces a new method that could guide treatment options for antibiotic-resistant UTIs, aiming to lessen antibiotic use by combining the pathogens’ unique metabolic traits.

Graphical Abstract
Metatranscriptome sequencing was used to investigate the functional uromicrobiome across a cohort of 19 individuals; patient-specific microbiome community models were reconstructed and simulated in a virtual urine environment. Total RNA was extracted from patients’ urine and sequenced to assess the metatranscriptome, providing insights into patient-specific uromicrobiome microbial taxa and their associated gene expression during urinary tract infections (UTIs). These combinatory datasets derived from metatranscriptomics data were further expanded first to reconstruct species specific metabolic models that were conditioned with gene expression. Gene expression conditioned metabolic models were combined in an in silico environment with a defined urine media to construct patient-specific context-specific uromicrobiome models, enabling an understanding of each patient’s unique microbiome. Using this approach, we aimed to identify patient-specific microbiome dynamics and provide insight towards various metabolic features that can be utilized or validated in future studies for individualized intervention strategies. Created with www.biorender.com.

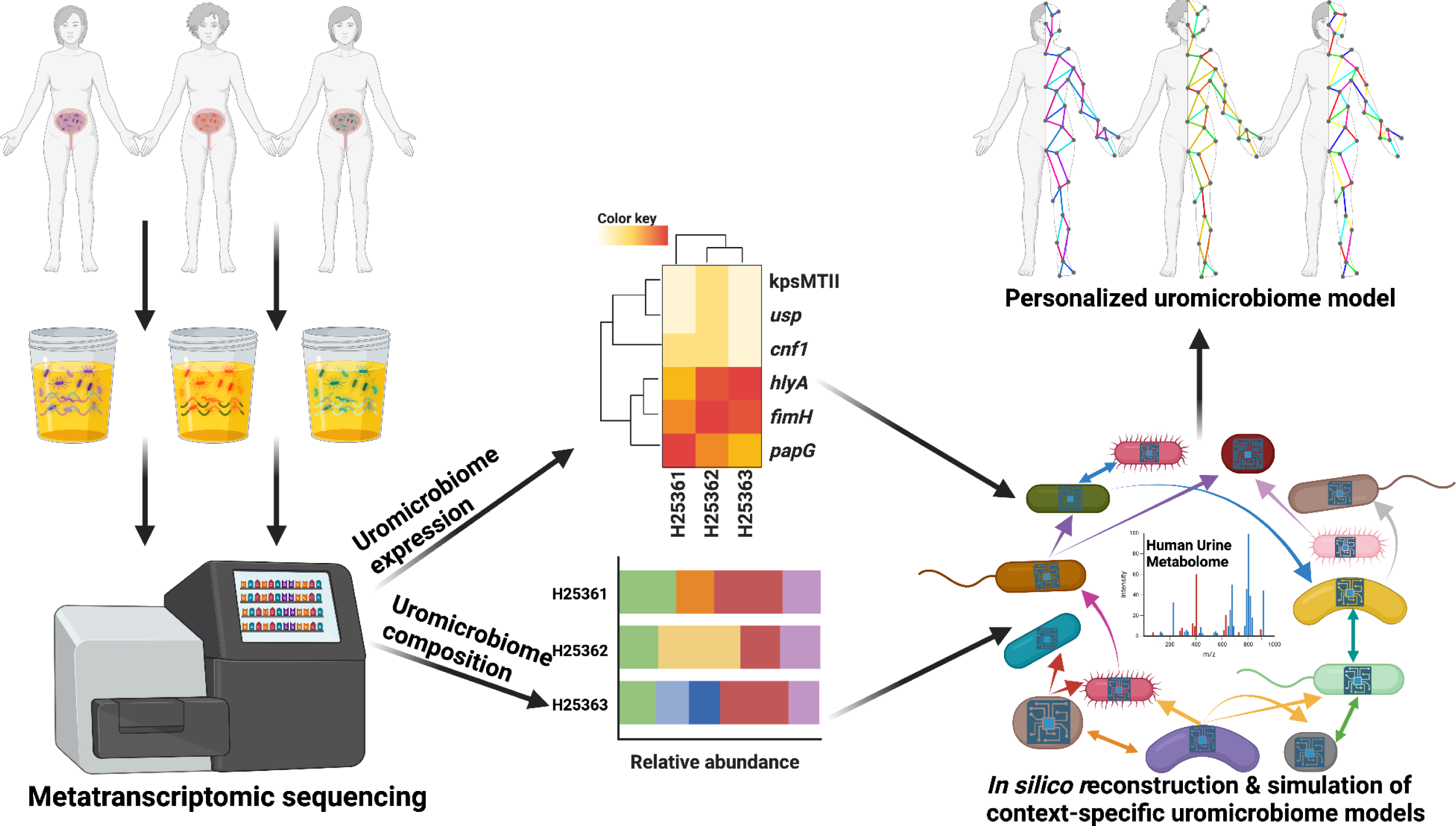

## I. Introduction

Urinary tract infections (UTIs) persist as one of the most common global health problems, with significant annual cases and healthcare costs reported worldwide (Flores-Mireles et al. 2015). Specifically, uncomplicated UTIs not associated with anatomical abnormalities frequently affect women and may result in recurrent, chronic infections (Hooton, 2012). UTIs are diagnosed by the presence of uropathogenic *E. coli (*UPEC), a bacterium found in approximately 80% of UTI cases (Flores-Mireles et al., 2015). Even with identifying a primary causative factor, the growing issue of multi-drug resistant UTIs has compounded effective treatment; present antibiotic treatments have become increasingly ineffective and led to the elevation of UTIs from common infections to intricate medical challenges (Mediavilla et al. 2016; Shields et al. 2021). As a result, the need for a detailed understanding of UTI-associated microbial ecology and functional consequences of metabolic crosstalk between pathogen communities in humans has become pressing to develop novel therapeutic approaches for UTI treatment.

Scrutinizing microbial communities through shotgun metagenomics (sequencing of DNA from microbial communities) is routinely applied to investigate human diseases, but fewer studies aim to leverage metatranscriptomes (sequencing of RNA from microbial communities) across human niches. While metatranscriptomics is limited by low resolution in distinguishing species-level organisms from one another compared to metagenomics. However, metatranscriptomics offers the unique advantage that it can identify both microbial taxa and active transcripts which can be used to describe microbial community responses to human hosts (Ojala et al. 2023). For example, metatranscriptomics has been used to investigate the vaginal microbiome during vaginosis to study *G. vaginalis* and how this microbe circumvents stress from the antibiotic metronidazole by expressing genes for DNA repair (Deng et al. 2018). While RNA-seq alone has been utilized to investigate how uropathogenic *E. coli* responds in human urine and activates virulence mechanisms (Hagan et al. 2010), RNA-seq focuses only on species-specific responses. At the same time, metatranscriptomics can investigate the gene response from entire human-associated microbial communities.

To expand the utility of sequencing human microbiomes through mechanistic studies of community interactions and phenotypes, the application of genome-scale metabolic models (GEMs) is becoming widely popular. GEMs are mathematical reconstructions of the metabolic network of a species, this approach provides a consolidated computational view into organismal physiology (O’Brien et al. 2015). An increasingly common and robust approach to investigating microbial response to specific conditions or contexts is the application of GEMs constrained by OMICs data (transcriptomics, proteomics, or metabolomics) to model microbial response to their environment effectively (Gopalakrishnan et al. 2023, Sen et al. 2023). While the AGORA2 database has provided a resource of 7203 gut-derived GEMs, which can be used to model drug transformation or bioremediation in a human niche accurately, context-specific GEMs have also been applied to study whole-body metabolism to simulate the interaction between hosts and microbes (Thiele et al. 2020; Heinken et al 2023). Recently, a reproducible and flexible framework has been developed to combine metatranscriptomics with GEMs to improve the functional understanding of microbial communities (Zampieri et al. 2023).

This study employs a systems biology approach to explore UTI pathology and microbial community physiology by integrating metatranscriptomics with metabolic modeling to evaluate microbial functions from clinical isolates. We analyzed 19 UTI patient samples and assessed their respective microbial diversity and gene expression to construct GEMs for uropathogen communities to predict metabolism in these human-associated ecosystems. Simulations of microbial growth and cross-talk were enhanced by applying a generic *in silico* urine medium based on the comprehensive Human Urine Metabolome database (Bouatra et al. 2013; Noronha et al. 2018), aiding in reconstructing patient-specific uromicrobiomes. Our findings offer insights into the variable microbial responses in UTIs, supporting the development of personalized treatments which rely on metabolic reprogramming, rather than antibiotic treatments.

## II. Material & Methods

### linical assessment and study criteria

Samples used to reconstruct and analyze microbiome community models herein are derived from a study cohort of patients sampled from November 2018 to August 2019 at the University Hospital Schleswig-Holstein and the University of Lübeck. The study was approved by the institutional review board (“Ethikkommission”) of the University of Lübeck on February 1st 2018 with the application number AZ18-009. The study comprises female patients with acute symptoms of uncomplicated UTIs who had confirmed evidence of uropathogenic *E. coli* that were cultured from self-sampled urine. We confirmed infection with *E. coli* by incubation of fresh urine on chromID CPS Elite agar plates (Biomérieux SA, Marcy-l‘Étoile, France). If *E. coli*-like appearance of growing colonies was identified on the plates, we further classified colonies using a Maldi Biotyper (Bruker Daltonik GmbH, Bremen, Germany) and included samples into the study if results were clearly interpretable as *E. coli* in two independent measurements. All participants were 18 years or older at the time of the study and sampling was performed before antibiotic therapy was initiated. The study excluded patients who exhibited physical malformations or comorbidities indicative of a complicated UTI. These include anatomical urogenital malformations (urethral stricture, bladder stones or tumors), previous urinary tract surgery, neurological or other diseases affecting bladder emptying, prior slight pelvis radiation, cytostatic treatment within the past year or during the study, immunosuppression, type 1 or 2 diabetes mellitus, pregnancy, recent use of vaginal probiotic therapies or vaginal spermicides, and participation in clinical studies potentially affecting urinary or renal function within 30 days before or during the study.

### Nucleic Acid Extraction from Urine Samples

#### RNA Isolation and Metatranscriptomic Sequencing

Total RNA was extracted from urine samples using the MICROBEnrich and MICROBExpress mRNA enrichment kits (Ambion, Thermo Fisher Scientific), which selectively reduce human RNA contamination and isolate bacterial RNA. RNA purification was performed using the TruSeq Stranded Total RNA kit (Illumina) to enable shotgun metatranscriptomic sequencing and facilitate functional profiling of the urinary microbiome. Metatranscriptomic sequencing was conducted at the Competence Centre for Genomic Analysis (CCGA) at Christian-Albrecht University, Kiel. Initially, samples H25361–H25365 were sequenced on an Illumina MiSeq platform, generating 2×75 bp paired-end reads. As additional resources became available, 14 more samples were sequenced on an Illumina NovaSeq 6000 platform with 2×50 bp paired-end reads.

#### DNA Isolation and 16S rDNA Gene Sequencing

DNA was isolated concurrently with RNA from fresh urine samples using the AllPrep DNA/RNA Mini Kit (Qiagen GmbH, Hilden, Germany), following the manufacturer’s protocol. To control for reagent contamination, negative isolation controls were included in every round of extraction and subjected to the same downstream processing as the samples. Extracted DNA was stored at −20°C for further analysis. For 16S rDNA gene sequencing, DNA was extracted from 19 metatranscriptomic samples, of which 15 were selected for targeted amplification of the V3/V4 hypervariable regions. Partial 16S rDNA gene sequences were amplified using primers containing linkers and indices designed for the V3/V4 regions, as described in Graspeuntner et al. (2018) and updated in Graspeuntner et al. (2024). Amplified products were pooled in equimolar concentrations, purified using the MinElute Gel Extraction Kit (Qiagen), and quantified before library preparation (Lüth et al. 2022). Sequencing was performed on an Illumina MiSeq platform using the MiSeq Reagent Kit V3 (600 cycles). The PhiX sequencing library was included as a positive control to monitor sequencing accuracy, and negative isolation controls remained negative throughout. Details on each sample, their identifying codes, and isolation source are in **Supplementary Table S1**.

#### Library Quality Controls and Sequencing Validations

For all sequencing workflows, rigorous quality controls were implemented. Negative isolation controls were processed alongside samples to ensure reagent purity and detect potential contamination. Only samples with defined amplicons after PCR amplification proceeded to sequencing and downstream data analysis, while controls confirmed the absence of contamination.

### Bioinformatics analysis and pre-processing microbial community models

#### Metatranscriptomics Preprocessing

An overview of the metatranscriptomics workflow, beginning with raw RNA reads and concluding with the reconstruction of context-specific microbiome community models, is provided in **Figure 1**. The preprocessing pipeline starts with quality control (QC) of both raw total RNA and 16S rDNA sequencing data. This step ensures high-quality reads, separates mRNA from rRNA in metatranscriptomic data for metabolic modeling, and isolates microbial rDNA, which serves as a positive control for filtering and validation. Sequence quality was assessed using fastQC (v0.11.8) (Wingett et al., 2018), identifying low-quality reads for removal both before and after QC. Low-quality and repetitive sequences were trimmed using Cutadapt (v1.5) (Martin, 2011) and PRINSEQ-lite (v0.20.4) (Schmieder et al., 2011). With Cutadapt, adapters and low-quality sequences were trimmed from paired-end reads. The minimum read length was set to 1, and reads with a quality score below 10 were removed. Overlap detection required a minimum of 5 matching bases. Several adapter sequences were specified for removal, including Illumina adapters and sample-specific contaminants. Reads that were input into PRINSEQ were filtered to retain a minimum length of 1 base and an average quality score of at least 10. Up to 5 ambiguous bases per read were allowed. Low-quality bases were trimmed from both ends, with up to 10 bases removed from the tails on the left and right.

**Figure 1:**
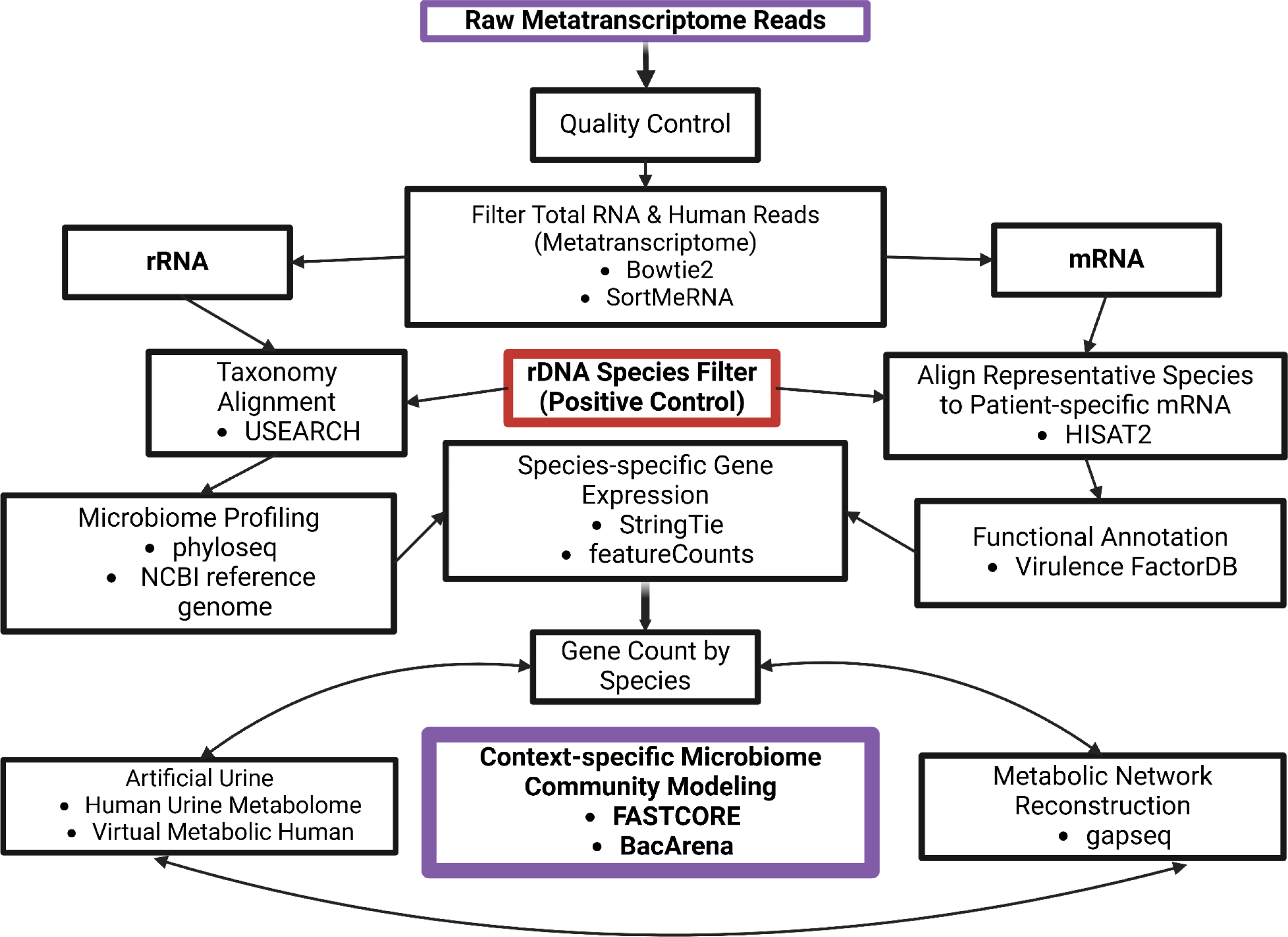
Workflow of context-specific microbiome community model reconstruction based on metatranscriptomic data. Our computational workflow processed 19 raw metatranscriptomic datasets into context-specific microbiome community models for exploring UTI-related metabolic interactions. After splitting the metatranscriptomic data into rRNA and mRNA fractions, two separate pipelines were used to study taxonomic and functional profiles, respectively, before combining them again to reconstruct context-specific metabolic models. Namely microbial rRNA was clustered into OTUs and aligned microbial databases, prior to microbiome profiling. Identified species were verified through complementary 16s rDNA sequencing. Reference genomes for the identified species were accessed from NCBI to map metatranscriptome-derived mRNA to the samples and in tandem, reconstruct metabolic models based on these reference genomes. Species-specific gene-level counts were mapped to reconstructed models to create context-specific metabolic models, which were representative of gene expression derived from patient uromicrobiomes. Patient-specific microbiome community members were added into a virtual simulation with a developed media from the Human Urine Metabolome to simulate context-specific uromicrobiome cross-feeding and environmental interactions.

To remove human RNA contamination, total RNA reads were aligned to a human reference genome using Bowtie2 (v2.5.4) (Langmead et al., 2012). Paired-end reads were mapped against the GRCh38 human reference genome (GCF_000001405.26) using default parameters for sensitive alignment. Aligned reads, representing human-associated sequences, were removed from the dataset, with only non-human reads being retained for downstream metatranscriptomic analysis. SortMeRNA (v4.3.7) (Kopylova et al. 2012) was used to classify the metatranscriptome, employing seven indexed databases, including the SILVA rRNA database (Yilmaz et al., 2014) and Rfam (Kalvari et al., 2021), for the identification of patient RNA. Databases such as Silva-Bac16s and Silva-Bac23s were used to extract bacterial rRNA. Non-mapped reads were presumed to be mRNA and retained for functional analysis and reconstruction of context-specific metabolic models. Mapping rates for each sample are detailed in **Supplementary Table S2**. All pre-processed reads (i.e rRNA and mRNA derived from metatranscriptomes and 16s rRNA derived from rDNA) were subject to CD-HIT (v4.8.1) (Fu et al. 2012) to reduce redundancy in the datasets by clustering highly similar sequences. For each sample, CD-HIT was executed with an identity threshold of 95% and a word size of 5.

### Metatranscriptomics Taxonomy and Species Filtering

To identify microbial taxa in both metatranscriptome-derived rRNA and 16s rDNA paired-end reads were merged in USEARCH(v12.0) (Edgar, 2010) with the ‘fastq_mergepairs’ command. To remove redundancy, the ‘fastx_uniques’ command was used to identify unique sequences and exclude singletons, to generate a non-redundant dataset where each sequence was labeled and size information was appended. Denoising and chimera filtering were performed with the ‘unoise3’ command, which clustered sequences into zero-radius OTUs (zOTUs) based on 95% sequence identity, to remove sequencing errors and chimeric artifacts. For zOTU clustering, a bootstrap confidence of 95% was applied, and sequences were screened to retain only biologically relevant OTUs. To quantify microbial abundance, the ‘usearch_global’ was used to match sequences to zOTUs with a minimum identity threshold of 95%. Finally, taxonomic classification was performed using the ‘sintax’ command against the RDP 16S reference database, with a cutoff of 0.95 for confident taxonomic assignments. Post-QC, mRNA and rRNA reads were categorized into transcriptional units (TUs) and operational taxonomic units (OTUs) using the USEARCH(v12.0) algorithm (Edgar, 2010) and plotted with a PCA to identify any sources of technical bias from differences in sequencing technology (**Supplementary Figure S1**).

Phyloseq (v1.5) (McMurdie et al., 2013) was used to assign taxonomy and estimate the relative abundance of microbial communities within each patient sample. By aligning zOTU and taxonomy tables derived from USEARCH with manually curated sample metadata, a *phyloseq* object was created. Low-abundance OTUs were removed from the downstream analysis if OTUs were present in fewer than two samples or with fewer than ten reads. Relative abundance normalization was applied using the ‘transform_sample_counts’ function which divided OTU counts by the total number of reads in each sample. Strains with sufficient representation across samples were retained to include species with at least one non-zero value above this level. Finally, to differentiate environmental sequences from host-associated microbial species found in the metatranscriptome reads, the 16S rDNA was used as a positive control filter. rRNA sequences derived from the metatranscriptomic dataset were cross-referenced with the 16S rDNA dataset. Only taxa present in both datasets were retained for downstream analysis. Species detected exclusively in the metatranscriptomic rRNA data but absent from the 16S rDNA dataset were excluded as potential environmental contaminants.

### Metatranscriptomic Species-specific Gene-level Counts

The remaining mRNA, separated by SortMeRNA, was used for functional analysis and species-specific gene expression quantification to inform community metabolic modeling. HISAT2 (v2.1.0) (Kim et al., 2019), SamTools (v1.16) (Danecek et al., 2021), and FeatureCounts (v2.0.3) (Liao et al., 2014) were used sequentially for mRNA mapping, alignment, and gene counting. Reference genomes and their annotations accessed from NCBI based on species identified from our prior taxonomic analyses (**Supplementary Table S3**), and were used to construct species-specific genome indexes with HISAT2. For each species, mRNA reads were aligned to the corresponding genome index using HISAT2 mode, producing SAM alignment files. Aligned reads were processed using SamTools to convert SAM files into binary BAM format followed by sorting and indexing. Sorted BAM files were subjected to FeatureCounts to assign reads to genes from genome annotations. StringTie (Pertea et al., 2015) was used to quantify gene expression and generate species-level transcriptomes. Sorted BAM files were used with genome annotations to calculate gene expression values, normalized as FPKM across samples. FPKM data were compiled into a master abundance table for comprehensive cohort-level comparisons. Given that all samples contained the *E. coli,* causative agent of UTIs, gene-level FPKM values were compared to the UPEC UTI89 reference genome (GCF_000013265.1) to identify virulence-associated genes and annotated with virulence factors from Virulence Factor Database (VFDB) (Liu et al., 2022). Virulence associated genes were categorized (Adherence, effector-delivery system, exotoxin, and nutritional/metabolic factor) based on their functional roles in pathogenicity to identify potential trends related to *E. coli UTI89* that have the potential to reveal clinically relevant pathogenic features.

### Reconstructing microbiome community models

#### *In silico* human urine medium

To enhance the accuracy and refinement of context-specific microbial community models, we created an *in silico* urine metabolic medium based on data from the Urine Metabolome Database (Bouatra et al., 2013), which cataloged 445 metabolites in healthy human urine via methods like NMR and GC-MS. From this pool, only 140 metabolites were found to be usable after cross-checking with matching naming and usability with the Virtual Metabolic Human (VMH) database as microbial exchange reactions (for example: if the metabolite common name is ‘phosphate’, the metabolic reaction will be listed as ‘Human (FALSE)/Microbe(TRUE)’ and will be noted with the exchange reaction ‘EX_pi(e)’), enabling the simulation of metabolite uptake and release (Noronha et al., 2019). We extracted and converted the provided metabolite concentrations from the Human Metabolome Database tables to millimolar units, normalized all values based on the identified average creatinine levels across the database, and ensured all acids were in their conjugate base forms (Marinos et al. 2020). A table which contains the output of this data processing of a *in silico* urine medium derived from the Human Urine Metabolome which includes metabolite details, exchange reactions, and concentrations for simulations, is detailed in **Supplementary Table S4**.

#### Reconstruction & preprocessing of context-specific metabolic models

Our workflow used the *gapseq* (v1.3.1) approach to reconstruct patient-specific GEMs for simulating microbial communities *in silico* (Zimmermann et al., 2021). Reference genomes of bacteria that were identified in the metatranscriptomics analysis and listed in **Supplementary Table S3** were used with *gapseq* to predict metabolic pathways, transport reactions, and associated metabolic networks. While this process was useful in reconstructing species relevant GEMs, these models were further refined with flux data derived from our developed *in silico* urine media that was sourced from the Human Urine Metabolome to mimic physiological conditions (**Supplementary Table S4**). To predict metabolic reactions, pathways, and transport systems associated with each species’ genome, the *gapseq ‘*find’ command was used with the ‘-p all’ option to include all pathway databases and the ‘-m bacteria’ flag to specify bacterial genomes with a BLAST bit-score threshold 200. The gapseq ‘find-transport’ command identified specific transport reactions using curated rules for membrane transport, needed to accurately model both nutrient uptake and metabolite exchange. A draft metabolic model for each species was then created using the *gapseq ‘*draft’ command, which combined the reaction, transport, and pathway data with the corresponding genome sequence. Parameters included upper bounds on metabolic flux, pathway weightings, and simulation constraints such as stoichiometric and thermodynamic limits. This draft model was refined through a gap-filling step, which incorporated additional reactions necessary to achieve metabolic functionality in the context of our custom *in silico* urine medium. Gap-filling used a reaction weight table, a reaction-to-gene mapping file, and a predefined nutrient file (**Supplementary Table S4**) to optimize metabolic exchange pathways specific to the medium.

To generate context-specific metabolic models, we integrated gene expression data with species-specific GEMs derived using *gapseq*. First, gene expression was linked to metabolic reactions by mapping gene-level expression data normalized to FPKM values (from previously described transcript quantification pipeline) to genes within the GEMs based on chromosomal positions that were subsequently associated with metabolic reactions. The integration of these data with GEMs used the COBRA Toolbox (v2.45.2) in MATLAB (Heirendt et al. 2019). Expression data that was specific to species, from each patient sample was loaded, and reactions in the GEMs were assigned expression values using the ‘mapExpressionToReactions’ function. Expression values for reactions were computed as the minimum expression level among associated genes to ensure a conservative estimate. To identify an active metabolic reaction set, we applied the ‘fastcc’ function from the *fastcore* (Vlassis et al. 2014) suite to establish a consistent metabolic network. This step systematically removed inconsistent reactions that were not flux-capable under patient-specific conditions, refining the GEMs for stability in downstream simulations. Next, the *fastcore* algorithm (Vlassis et al. 2014) was used to reconstruct context-specific models by extracting core reactions from the consistent network based on expression thresholds, with reactions in the top 25% of expression values manually designated as “active” as suggested by Richelle et al. (2019a), the final reconstructed models were saved for further analysis. Flux balance analysis (FBA) was performed on each context-specific model following reconstruction to assess their metabolic capabilities to carry a consistent metabolic flux across their network. Biomass production was used as the objective function, and optimization was conducted using the ‘optimizeCbModel’ command. Successful FBA results confirmed the viability of the reconstructed networks.

### Simulation of microbiome-specific community models

Patient-specific community models were created by using the aforementioned reconstructed and validated context-specific GEMs in BacArena(v1.8.1) (Bauer & Zimmermann et al., 2017), by adding all patient-specific GEMs into a constrained arena to explore community growth, metabolic flux, and cross-feeding interactions through community FBA. Two simulation types were conducted on microbial communities related to patient-specific UTIs: 1) ‘non-context’ simulations, involving *gapseq* models gapfilled with the previously described i*n silico* urine medium and made consistent with *fastcc*, and 2) ‘context-specific’ simulations, which are the same as the previously described models, but with the addition of mapped gene expression data derived from metatranscriptomics to further constrain the models. The simulations aimed to uncover whether context-specific modeling as an experimental condition could reveal metabolic differences associated with pathogenicity. BacArena simulations were structured and implemented into R (v4.3.2) to investigate the community FBA of each patient microbiome as constrained by metatranscriptomics. First, the previously described artificial urine medium prepared derived from Human Urine Metabolome which contained extracellular exchange reactions from VMH and their respective flux values, were formatted to align with BacArena’s requirements. Patient-specific microbial related GEMs that constitute the host microbiome were read into R and assigned unique identifiers based on the species names. These GEMs were integrated into a 100×100 arena, simulating microbial communities with relative abundances derived from patient-specific datasets. Each species was added with its relative abundance, normalized against the total community abundance. The arena was initialized with 1000 organisms, distributed evenly to maintain substrate availability and organismal balance. Species diversity ranged across models, incorporating core pathogens like *E. coli UTI89* alongside other commensals. Urine-specific metabolites were added to the simulation as extracellular substances.Simulations were conducted over four iterations, each representing one hour, and repeated in triplicate. Outputs were analyzed to assess growth patterns, cross-feeding interactions, and metabolic outputs across all species. Metabolite utilization and production were extracted using functions like ‘findFeeding3’ to investigate statistical differences of community-level and species-specific fluxes to highlight metabolic behaviors and interspecies interactions within patient-specific urinary microbiomes.

### Analysis of community simulations

Predicted metabolic subsystem abundance data for each patient and their respective context-specific and non-context-specific microbiome community models were generated through BacArena simulations. To compare the statistical impact of applying context-specific metabolic modeling between experimental conditions, a paired t-test was performed for each subsystem to determine statistically significant differences in metabolic activity between conditions. P-values were adjusted using the Benjamini-Hochberg method to identify context-specific metabolic subsystems with adjusted p-values below a significance threshold of 0.05. Significant resulting p-values were transformed to their negative logarithmic form (-log10). We focused our comparison of context-specific versus non-context-specific metabolic functions by studying metabolic flux distributions and their activity patterns across both experimental conditions and microbial communities. Samples are classified into three manually defined microbial community types—*Lactobacillus Diverse* (more than three species (n=5)), *Lactobacillus Absent* (absence of Lactobacillus (n=10)), and *Lactobacillus Single* (dominated by a single species (n=4))—to evaluate their distinct impacts on metabolic activity. Statistical testing was applied between these three predefined groups by performing an ANOVA with Tukey’s post-hoc tests for pairwise comparisons (e.g., *Lactobacillus* Diverse vs. *Lactobacillus* Absent and *Lactobacillus* Diverse vs. *Lactobacillus* Single). Next, community wide metabolic flux data was then analyzed through fold-change patterns between experimental conditions for specific patients. Log fold changes are calculated for each metabolic subsystem by comparing activity levels under “Context” and “Non-context” conditions and low variance across samples are filtered out, retaining only those with significant variability above a defined threshold (0.30) to highlight relative differences between subsystems and experimental conditions. Hierarchical clustering was performed using Euclidean distance.

Metabolic fluxes in microbial communities, specifically focusing on both positive (production) and negative (consumption) fluxes across various reactions and metabolites. Using predictions of metabolic flux measurements from BacArena simulations we identified positive fluxes greater than zero, indicating metabolic production processes, and negative fluxes less than zero, indicating metabolic consumption processes. Each flux type was analyzed with a one-sample t-test to assess whether the flux deviates significantly from zero. The top reactions for both production and consumption are selected based on statistical significance and magnitude, with reactions exhibiting the highest mean flux values being prioritized. Finally, to investigate metabolic cross-feeding in microbial communities, we expanded our analysis of metabolite flux data and microbial production (positive flux) and consumption (negative flux) rates across organisms for various metabolites. The focus was on extracellular exchange (EX_) reactions, from which average fluxes were calculated for each metabolite-organism combination to identify the most actively exchanged metabolites within the community to reveal metabolic dependencies between species.

## III. Results

### UTI pathogenic microbiome composition and diversity vary across patients

Our analysis revealed diverse UTI bacterial communities among patients, indicating inter-patient biodiversity variation in terms of composition and abundance (**Figure 2**). **Figure 2A** presented species-level relative abundance, showing a range of identified microbial species that define a microbial signature for each patient. Variable abundance of microbial groups were observed, including genera such as *Anaeroglobus, Barnesiella, Blautia, Dialister, Escherichia/Shigella, Lactobacillus, Peptoniphilus, Porphyromonas, and Prevotella* (**Supplementary Figure S2**). Interestingly, while this analysis identified genera that are known to be potential urinary tract pathogens, it was noted that these human pathogens are generally associated with the gut, oral, and vaginal-associated environments were also observed; these pathogens may have roles in urinary tract colonization and warrant further investigation (Tufon et al. 2020; George et al. 2024; Miller et al. 2024; Pan et al. 2024; Russo et al. 2024; van Teijlingen et al. 2024). *Lactobacillus* species were prevalent, with patients having anywhere between 0-5 different species per sample; a high abundance and diversity of *Lactobacillus* species were observed in samples A01, B02, and G01. The impact of the abundance of these probiotic strains can be distinguished in the Principal Component Analysis (PCA) of species composition which underlies that diversity in the studied uromicrobiome’s varied ecological landscape, with PC1 and PC2 explaining 38.03% and 25.30% of the variance, respectively (**Figure 2B**). Shannon’s Alpha Diversity analysis presented a diversity range from 0.064 to 1.962 (**Figure 2C**), indicating a variability in this already simplistic community. Finally, a map of all species that were found across the samples UTI cohort in relative abundance, per patient was reconstructed to inform the downstream selection of species for inoculation into our microbial community modeling simulations, with a range of 5 to 15 species identified depending on the patient (**Figure 2D**).

**Figure 2:**
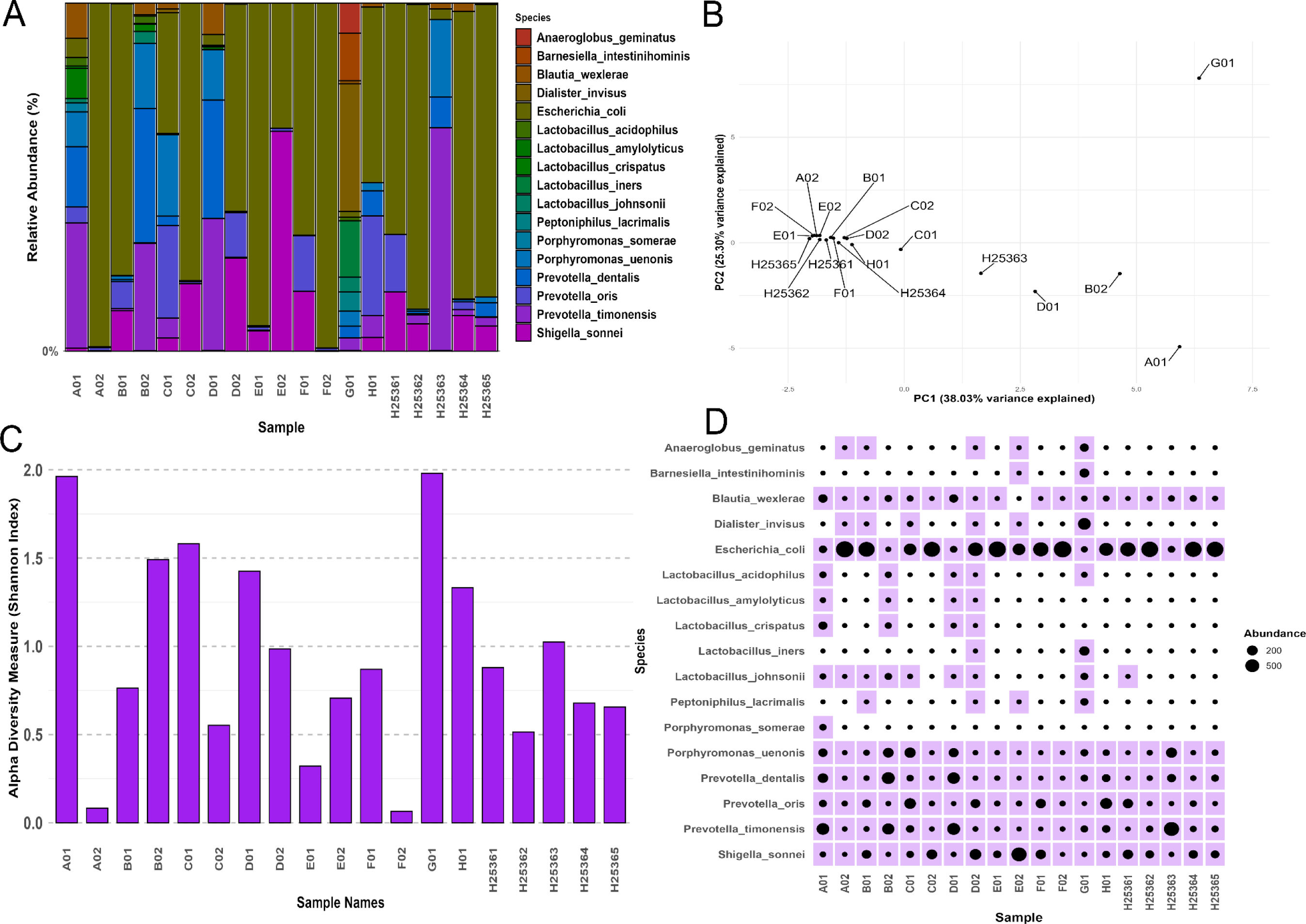
Diversity and Composition of UTI Microbiomes Across Patients. (A) Species-level relative abundance profiles present a variety of microbiome compositions across patients, with several known uropathogenic and non-urogenital taxa, including Anaeroglobus, Barnesiella, Escherichia/Shigella, and Lactobacillus. Notably, gut, oral, and vaginal-associated genera were also identified, highlighting the complex nature of urinary tract colonization and the need to expand present understanding on the crosstalk between host-associated microbiomes. (B) Principal Component Analysis (PCA) of species composition, with PC1 and PC2 explaining 38.03% and 25.30% of the variance, respectively. The PCA plot reveals substantial variability across patient microbiomes. While most patient groups cluster together, outliers that are likened to a high diversity of Lactobacillus species in samples A01, B02, and G01. (C) Shannon’s Alpha Diversity analysis showing diversity indices ranging from 0.064 to 1.962, indicating significant variability in microbiome complexity across the patient cohort. (D) Relative abundance map of species detected across all patient samples, with 5 to 15 species identified per patient. This map informs species selection for downstream microbial community modeling and simulations.

### UPEC virulence strategies are diverse across UTI patients

Gene expression was mapped to the UPEC UTI89 reference genome to extract FPKM value, focusing on the uropathogenic *E. coli* strain that epitomizes the primary cause of about 80% of UTIs (Flores-Mireles et al., 2015). The top-10 expressed genes for UPEC UTI89, mapped across multiple patient samples, revealed both conservative and distinct patterns of transcriptional activity as shown in **Figure 3A**. Across all samples, *ssrA* emerged as the most consistently abundant transcript. While other frequently detected transcripts were observed across all groups, including *rnpB, cspA*, and *ssrS*. Most interesting, is the observation of several highly expressed UTI89 locus tags which have unknown annotations and functions in relation to uropathogenic virulence.

**Figure 3:**
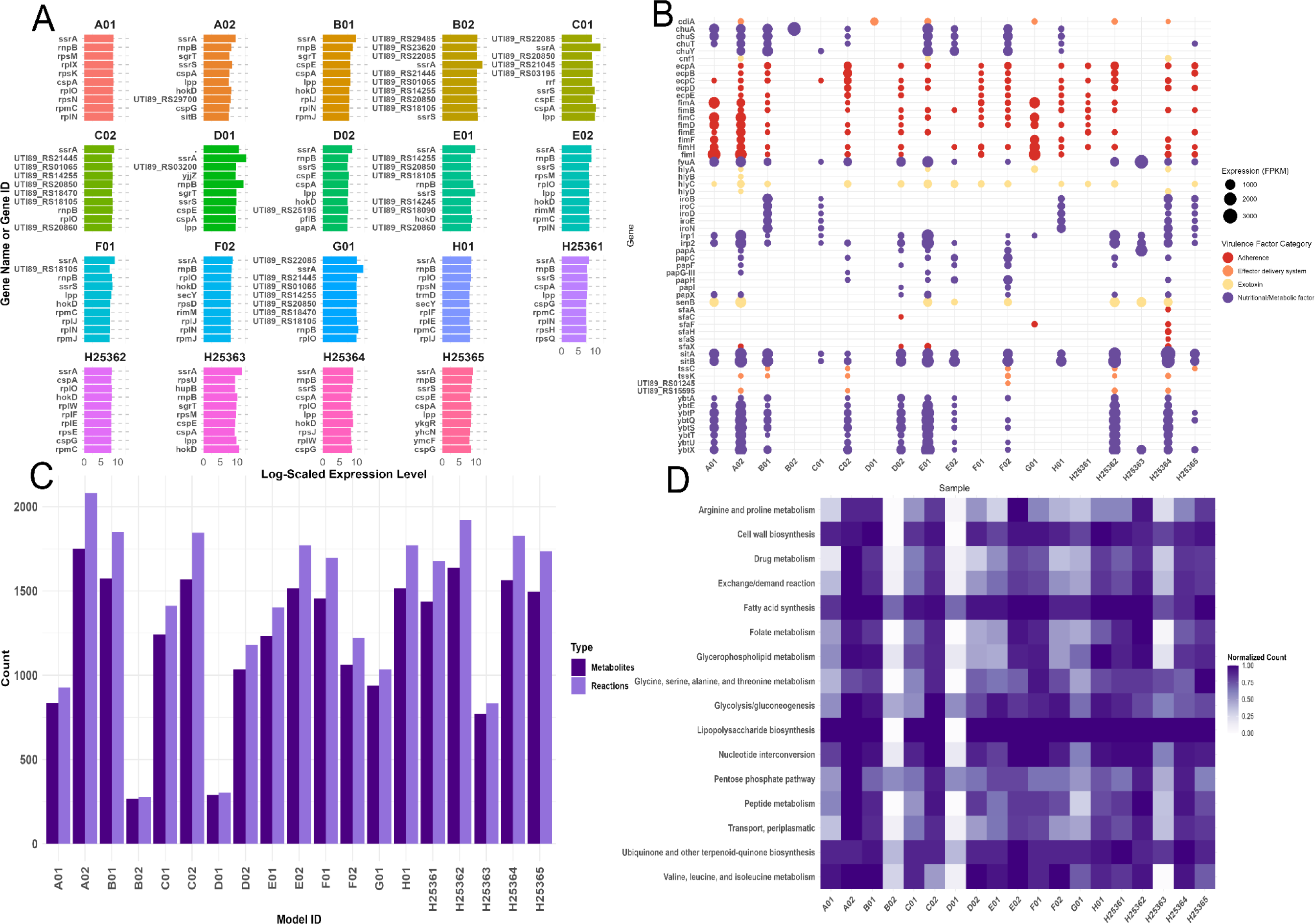
Diverse Virulence Strategies and Metabolic Adaptation in UPEC UTI89 Across Patients. (A) Gene expression profiles were extracted for the top 10 most abundant transcripts in UPEC UTI89 across multiple patient samples. Profiles were normalized through log-scaled expression levels to assess virulence strategies across patients. The results presented that the gene ssrA was consistently expressed across all samples, while other genes, such as rnpB, cspA, and ssrS, presented more variable expression patterns throughout the compendium of UPEC UTI89 strains. (B) Clinically relevant virulence factors were further assessed based on strain-specific gene expression from each patient, and linked to known pathogenic mechanisms of UPEC UTI89 using the Virulence Factor Database. Specifically, the analysis presents high expression levels of adhesion-related genes (fimA and fimI) and iron uptake genes (chuY, chuS, iroN) across most patient samples. These findings are suggestive of a consistent signature of UPEC UTI89 adaptation strategies that can be observed during UTI infection and host colonization. (C) Reconstructed context-specific metabolic models of patient-specific UPEC UTI89 strains presented variations in metabolic diversity. For example, strains exhibiting varying numbers of reactions and metabolites, ranging from over 2,000 reactions and 1,800 metabolites to fewer than 300. (D) Metabolic subsystem activity profiles across patient-specific UPEC UTI89 strains derived from context-specific metabolic models. Despite conserved gene expression, diverse variability was observed in the normalized count activity of metabolic subsystems within the metabolic model, including Arginine and proline metabolism, Glycolysis/gluconeogenesis, Peptide metabolism, and Valine, leucine, and isoleucine metabolism which can be likened to UPEC strains’ complex approaches to resource utilization in host environments.

Gene expression signatures were further associated with various virulence factor traits by cross-referencing gene expression to common etiological factors of these infections. By annotating gene expression profiles with functional data from the Virulence Factor Database (Liu et al., 2022), clinically relevant pathogenic mechanisms of UPEC UTI89 within our cohort were revealed. Virulence genes with high expression in patient samples highlighted the diversity of virulence strategies (**Figure 3B**). Specifically, *fimA* and *fimI* genes, important factors for bacterial adhesion, were expressed across various samples, emphasizing their classical role in establishing infections in host epithelial cells (Baker et al., 2021). Nutrient acquisition genes that are associated to iron uptake, such as *chuY*, *chuS*, and *iroN* were also observed across multiple samples, highlighting another common virulence mechanism that is leveraged by UPEC to adapt and invading host environments through metabolic modification (Mey et al. 2021; Edison et al., 2023; Zhou et al. 2023). This variability in clinically-relevant virulence factor gene expression underscores the flexibility of UPEC pathogenicity and suggests a range of clinical manifestations across different patient-specific infections, underlining the complexity of UPEC UTI89 pathogenic mechanisms and their potential to colonize diverse host environments.

Subsequently, we reconstructed, context-specific metabolic models of patient-specific UPEC UTI89 as part of our microbiome community modeling workflow, and specifically scrutinized to identify any metabolism related patterns. While some gene expression patterns from the previously described analysis were conserved across strains, mapping whole gene expression signatures to metabolic networks revealed distinct differences between these apparently related strains. The number of reactions and metabolites varied widely across strains, with some presenting greater metabolic diversity (**Figure 3C**). For example, certain strains exhibited over 2,000 reactions and nearly 1,800 metabolites within their metabolic models, suggesting a highly diverse metabolic network, while others had fewer than 300 reactions and metabolites, indicating a more constrained or conservative metabolic capacity.

The previously mentioned reconstructed UPEC UTI89 context-specific models were further investigated to reveal trends across the diversity in metabolic subsystem activities (**Figure 3D**). For instance, subsystems like ‘Arginine and proline metabolism’ presented variation in activity across our study, with some strains (A02) exhibiting high normalized counts (0.882), while others (B02 & D01) displayed little or no activity. Similarly, the ‘Drug metabolism’ and ‘Glycolysis/gluconeogenesis’ pathways were strongly enriched in strains derived from patients A02 and C02, but had lower activity in D01. The ‘Nucleotide interconversion subsystem’ was also variable, with strains derived from patients F02 and H01 presenting a high normalized activity, while strains from patient H25363 and H25365 having. With regards to the ‘Pentose phosphate pathway’, A02 presented maximal activity, whereas strains derived from patients D01 and B02 were largely inactive. Overall, these findings further highlight the need to move away from solely gene expression based assays and to consider the significant strain-specific variation in predicted metabolic networks. Specifically, these results suggest that UPEC’s metabolic capabilities are highly adaptable to specific host environments, and likely the key to its pathogenicity with regards to patient-specific interventions.

### Differential Microbiome Metabolic Activity in Context-Specific and Non-Context-Specific Models

We generated microbiome-wide metabolic subsystem abundance data for each patient to compare the metabolic differences between two experimental approaches: context-specific and non-context-specific microbiome community models, using BacArena simulations (**Figure 4A**). The two simulation types focused on microbial communities associated with patient-specific UTIs. First, *non-context-specific simulations* utilized gap-filled gapseq models based on the in silico urine medium, refined for consistency using fastcc. Second, *context-specific simulations* incorporated the same models but included additional constraints derived from metatranscriptomics, mapping gene expression data to further refine the models. These simulations aimed to evaluate whether integrating context-specific data could reveal distinct metabolic patterns linked to pathogenicity. Paired t-tests of subsystem activity presented significant statistical differences during the application of context-specific conditions, particularly in pathways such as ‘Biosynthesis of siderophore group nonribosomal peptides’ (log adj p-value = 3.14), ‘Fatty acid synthesis’ (log adj p-value = 3.01), and ‘Glycolysis/gluconeogenesis’ (log adj p-value = 3.01). Additional subsystems, including ‘Cell wall biosynthesis’ and ‘Peptide metabolism’ (both log adj p-value = 3.14), also showed significant alterations. The summed components of the PCA explained 30.30% of the variance (**Supplementary Figure S3A**), highlighting reduced variability in context-specific models compared to the outlier-prone non-context-specific models, which exhibited extraneous metabolic reactions. Context-specific models showed reduced total metabolic flux (**Supplementary Figure S3B**), suggesting that the application of context-specific modeling is more reflective of a focus on biologically relevant pathways. Euclidean distance analysis also presented sample-specific differences in metabolic flux, which is also indicative of distinct metabolic adaptations to environmental context when applying context-specific modeling approaches (**Supplementary Figure S3C)**. The presented results support the biological refinement in metabolic networks through the integration of context-specific data with the aim to improve the biological relevance and accuracy of microbial metabolic modeling.

**Figure 4:**
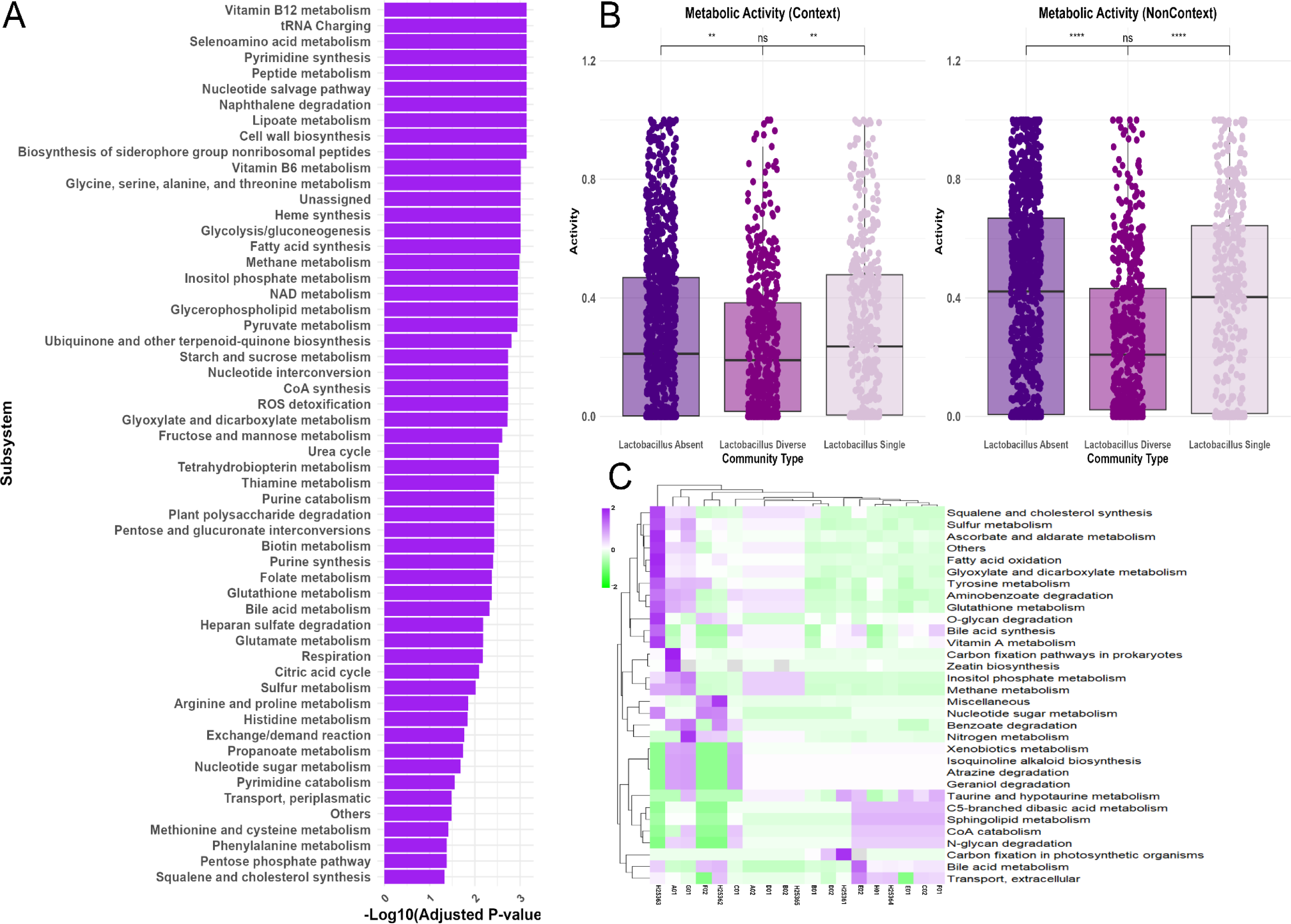
Differential Metabolic Activity Across Context-Specific and Non-Context Conditions in Uromicrobiomes. (A) Significant changes in metabolic subsystem activity were extracted from context-specific microbiome models in comparison to non-context-specific conditions. Metabolic subsystems such as ‘Biosynthesis of siderophore group nonribosomal peptides,’ ‘Fatty acid synthesis,’ and ‘Glycolysis/gluconeogenesis,’ were some of the most significant metabolic groupings, with adjusted p-values < 0.001. Paired t-tests were performed, and results were adjusted for multiple comparisons using the Benjamini-Hochberg method. (B) Given the diversity of observed Lactobacillus strains in the studied UTI patients, comparisons between the metabolic activities of three manually defined Lactobacillus community types were statistically assessed: Lactobacillus Diverse (multiple species (n=5)), Lactobacillus Single (dominated by one species (n=4)), and Lactobacillus Absent (no Lactobacillus present (n=10)). The Lactobacillus Absent group presented the highest mean metabolic activity in both context (0.272) and non-context (0.390) conditions, with significant differences (p < 0.001) between Lactobacillus Diverse and Lactobacillus Absent. Statistical comparisons were conducted using ANOVA followed by Tukey’s post-hoc tests. (C) A heatmap of log fold changes across metabolic subsystem activity between context-specific and non-context-specific conditions was useful to identify metabolic subsystems that were significantly variability across our studied cohort (log fold change > 0.30). O-glycan degradation and bile acid metabolism displayed particularly high variability in certain samples (e.g., F02, H25362, H25363).

Due to the observation that several patients contained a range of 0-5 *Lactobacillus* strains in differing abundances, we compared the metabolic functions between context-specific and non-context-specific microbial community models by examining the distribution and activity patterns of metabolic fluxes between our *in silico* experimental conditions (context-specific vs. non-context-specific) and manually defined *Lactobacillus* community types (**Figure 4B**). All investigated samples were classified into three distinct *Lactobacillus* community types: *Lactobacillus* Diverse (consisting of three or more *Lactobacillus* species(n=5)), *Lactobacillus* Absent (lack of *Lactobacillus species*(n=10)), and *Lactobacillus* Single (presence of a single *Lactobacillus* species(n=4)). Statistical comparisons between each group in a pair-wise approach identified that the *Lactobacillus* Absent group had the highest mean metabolic activity in both context and non-context conditions, followed by *Lactobacillus* Single group, and *Lactobacillus* Diverse. Statistical comparisons when using the ANOVA test, followed by Tukey’s post-hoc tests, presented significant differences between the *Lactobacillus* Diverse and Lactobacillus Absent (p < 0.001). The *Lactobacillus* Single group showed a significantly higher rate of metabolic activities as compared to *Lactobacillus* Diverse (p < 0.001) groups in both context and non-context conditions. These findings provide additional information to the context of UTIs, in that the presence and diversity of *Lactobacillus* species have a potentially substantial effect on microbiome community metabolic activity. Additionally, the comparison of metabolic activity between context-specific and non-context-specific models indicated that non-context conditions generally resulted in higher metabolic activities in comparison to the context-specific conditions (p < 0.001), regardless of the *Lactobacillus* group. Overall, these results which compare statistical differences in metabolic activities between *Lactobacillus* diverse groups have been useful to identify the impactful role of microbiome members, in addition to the application of context-specific modeling in defining microbiome metabolic activity in ecosystems.

Finally, community metabolic flux data was further assessed by calculating fold-changes between our two computational reconstruction conditions, for individual patients and subject to hierarchical clustering (**Figure 4C**). Log fold changes were first computed by comparing metabolic subsystem activity between context-specific and non-context specific conditions for each patient. Metabolic subsystems with low variance across all studied patient samples were filtered out to retain subsystems with the highest variability (log fold change > 0.30), with the aim to present groups with the most significant differences observed when applying context-specific modeling. Among these, O-glycan degradation was identified to be the metabolic subsystem with the highest variability in context-specific groups, with a noted enriched metabolic rate in samples F02, H25362, and H25363. Similarly, Bile acid metabolism also presented notable increases in samples F02, H25362, and H25362.

### Metabolic Flux Analysis of Production and Consumption Processes in Microbial Communities

We reconstructed microbiome-wide community models for each patient through metatranscriptomic sequencing and generated two datasets per patient to compare metabolic differences between two experimental approaches: context-specific and non-context-specific microbiome community models, using BacArena simulations. We then analyzed metabolic fluxes within microbial communities, focusing on both positive (production) and negative (consumption) reactions (**Figure 5A**). Using BacArena simulations, we identified reactions with significant deviations in metabolic flux values and selected the top reactions based on both the magnitude of their mean flux values and statistical significance. Among the positive flux reactions, we observed several key production processes, such as Ferredoxin, Cholate Transport via Bicarbonate Antiport, and Glycerol-3-phosphate acyltransferase, all exhibiting high and statistically significant fluxes (p-value < 0.05). These reactions suggest active utilization of likely host-associated metabolites that would be crucial for community survival. Conversely, the top negative flux reactions, indicating community consumption, included Hypoxanthine Exchange, Cholanate Exchange, and 5-dehydro-D-gluconate reductase. These identified reactions support the trend that the UTI microbial communities may specialize in nutrient uptake, especially for pyrimidine scavenging and sugar metabolism.

**Figure 5.**
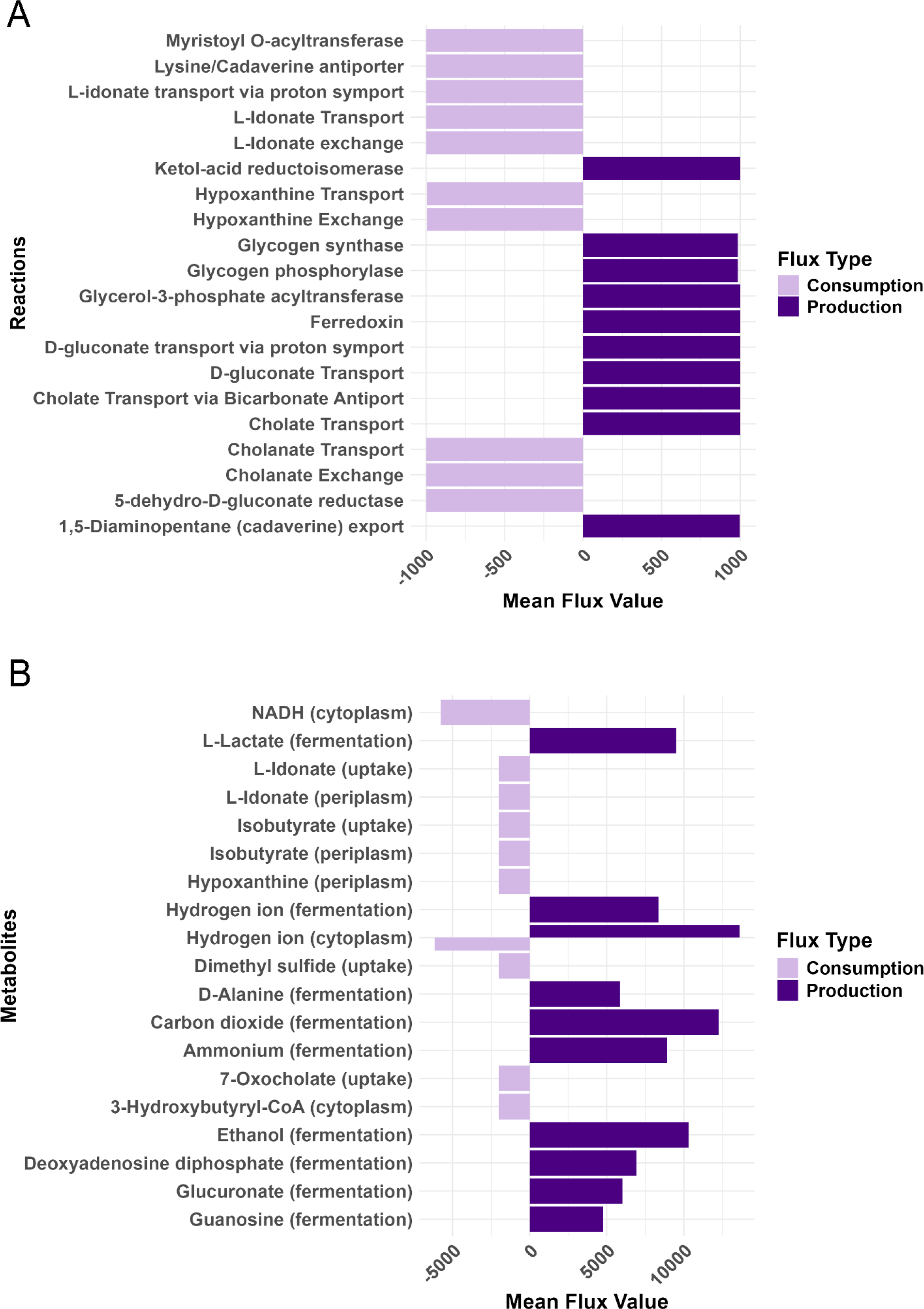
Metabolic Cross-talk and Product Contribution/Consumption in UTI Microbial Communities. (A) Analysis of metabolic fluxes within microbial communities of both positive (production) and negative (consumption) reactions. Top reactions were subset based on significant flux deviations through the combination of both the magnitude of mean flux values and statistical significance through a one-sample t-test (p-value < 0.05). Key production processes include Ferredoxin, Cholate Transport via Bicarbonate Antiport, and Glycerol-3-phosphate acyltransferase. Top consumption processes include Hypoxanthine Exchange, Cholanate Exchange, and 5-dehydro-D-gluconate reductase. Overall, the results from all microbial community models highlight the activity of metabolic reactions for uptaking nutrients for metabolic processes and the biosynthesis of essential products that are crucial for community survival. (B) Microbial community metabolites are subset in a similar statistical approach as previously mentioned to identify metabolites that are produced and consumed by microbial communities. Among the major metabolites produced via fermentation are Carbon dioxide, Ammonium, Hydrogen ion, Ethanol, and L-Lactate, that were all observed at significant levels. Other important metabolites of note are D-Alanine, Deoxyadenosine diphosphate, Glucuronate, and Guanosine were also produced. With regards to consumption of metabolites Hypoxanthine, 7-Oxocholate, and L-Idonate were notably uptaken, indicating a reliance on purine salvage and energy metabolism. The results from both analyses highlight a tightly regulated metabolic network that supports microbial community growth and adaptation through the usage of nucleobases and energy sources.

In addition to the major metabolites produced through fermentation, such as Carbon dioxide, Ammonium, Hydrogen, Ethanol, and L-Lactate, our analysis of uropathogenic community metabolism also identified several other microbial metabolic products (**Figure 5B**). For instance, D-Alanine, which is an important amino acid required for the biosynthesis of prokaryotic cell walls, was detected to be significantly produced when applying context-specific metabolic modeling, as assessed through a one-sample t-test, indicating the impact and importance of considering D-Alanine in clinical contexts to maintain pathogen cellular integrity and growth. Similarly, Deoxyadenosine diphosphate, Guanosine, and Hypoxanthine, were all observed to also be produced in high quantities, suggesting a high demand for purine biosynthesis and potentially cross-feeding within the microbial communities. In comparison to non-context-specific models, our t-test also identified Hypoxanthine, as an important nucleobase that was predicted to be consumed in context-specific community models, further highlighting a potential importance of purine salvage metabolism, exchange, and cross-feeding in the studied uropathogenic microbial communities. 7-Oxocholate (uptake), an intermediate involved in bile acid metabolism and cholate transport reactions, was also significantly consumed, further emphasizing the microbial community’s interactions with host-derived metabolites.Moreover, L-Idonate (periplasm) represents the metabolite’s extracellular availability, while L-Idonate (uptake) reflects its active transport into the cell, both contributing significantly to carbohydrate metabolism and cellular osmoregulation, supporting microbial homeostasis and energy acquisition in the host environment. Overall, the results reveal that the production and uptake of key metabolites is closely regulated and contributes to the diverse requirements of the core urogenital microbiome community’s survival, growth, and adaptation within the host environments.

### Interplay of metabolite cross-feeding in UTI microbial communities reveals potential pathogenic strategies

Metabolic cross-feeding across individual members of patient-specific microbiomes within microbial communities was investigated by analyzing metabolite flux data and examining the production (positive flux) and consumption (negative flux) rates of various metabolites between different organisms within the community. Our analysis focused specifically on metabolic exchange reactions, which are boundary reactions in metabolic networks that enable the import or export of metabolites between an organism and its environment, playing roles in nutrient uptake, waste secretion, and interspecies interactions. By calculating the average flux for each metabolite-organism combination, we identified trends in the most actively exchanged metabolites within the community **(Figure 6A**). Interestingly, several species of *Lactobacillus* which are commonly known for their probiotic properties, played a significant role in the production of Alpha-Lactose, a key metabolite involved in maintaining pH balance in the gut and vaginal microbiomes. *Lactobacillus* species, such as *L. rhamnosus* and *L. reuteri*, were also found to be involved in the fermentation of carbohydrates into short-chain fatty acids (SCFAs), which are important for maintaining gut health. In contrast, *UTI89* presented consumption of various SCFAs such as acetate, suggesting its role in nutrient acquisition and adaptation within the urinary tract. We also observed the unique production of ornithine by UTI89 and the consumption by the remaining members of the uromicrobiome, a metabolite which has been previously observed to provide a fitness advantage to pathogens. The interplay between *Lactobacillus* species and *UTI89* provides a unique example of microbial interactions, with *Lactobacillus* acting as a likely producer of metabolites that influence the growth and activity of pathogenic species like *UTI89*. However, direct cross-feeding of acetate between *Lactobacillus spp* and UTI89 was not observed, though it is possible that *Lactobacillus* is releasing acetate into the extracellular environment for UTI89 to uptake. Further analysis revealed other significant metabolic trends through a one-sample t-test within the community. Probiotic strains such as *L. amylolyticus* produced metabolites like L-threonine, hydrogen, and succinic acid, which are some of the most cross-exchanged metabolites in community metabolism. In contrast, *D. invisus* and *P. lacrimalis* showed consumption of metabolites like phosphate, L-glutamic acid, and D-glucose, highlighting their metabolic dependence on these compounds. Other organisms, such as *L. iners* and *B. intestinihominis*, demonstrated both production and consumption of a wide range of metabolites, such as taurine and glycine. These findings suggest dynamic interdependencies between species, with some microbes primarily acting as metabolite producers and others as consumers, facilitating complex cross-feeding interactions that influence community stability and function which shape the uromicrobiome ecosystem dynamics.

**Figure 6.**
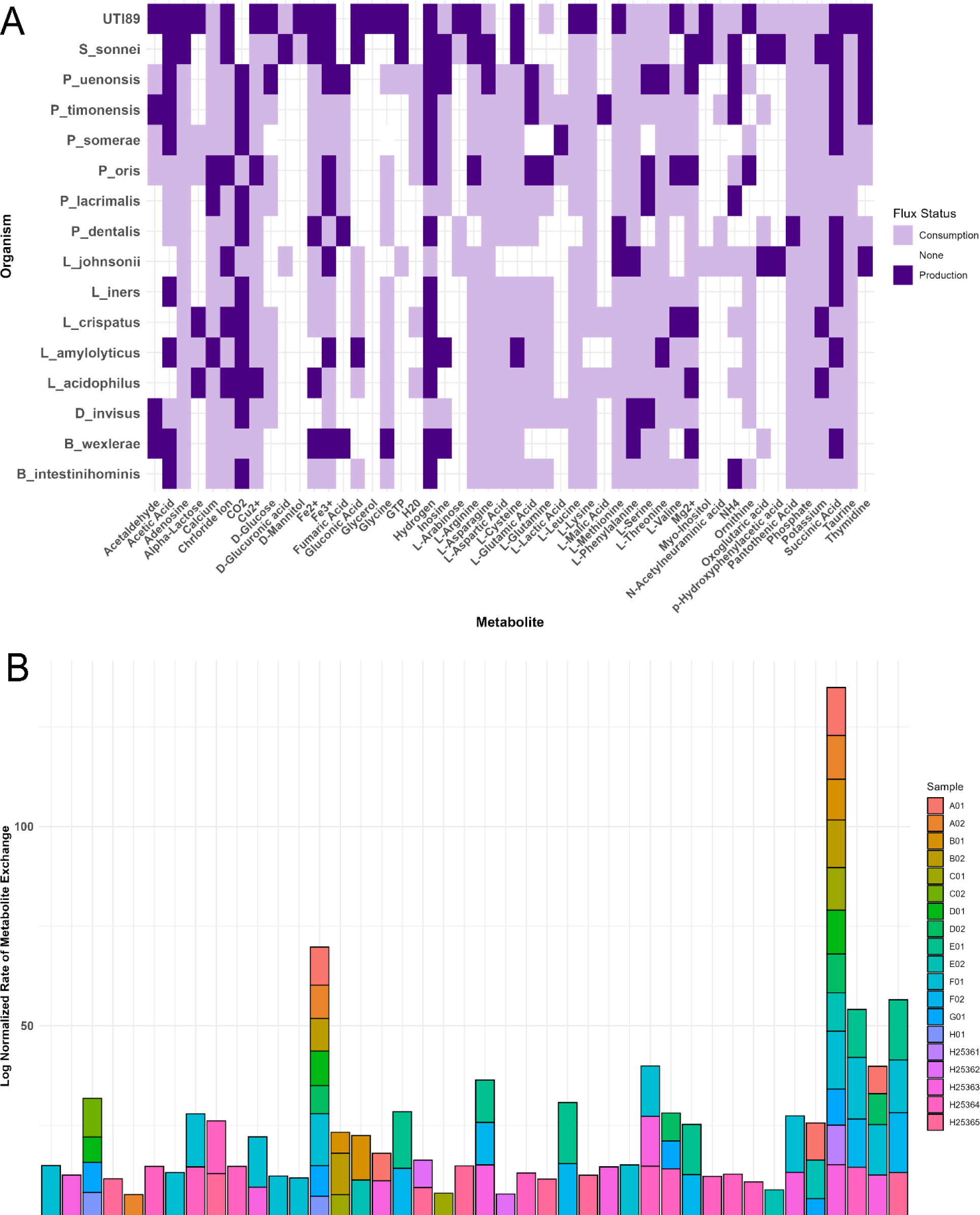
Analysis of Metabolic Cross-Feeding and Metabolite Exchange in Patient-Specific Microbiome Communities. (A) Metabolic cross-feeding within patient-specific microbiome communities was assessed through the production and consumption rates of various metabolites across microbial species. The analysis focused on extracellular exchange (EX_) reactions to identify important microbial contributors to metabolite exchange. (B) Log-normalized rates of metabolite exchange across the cohort of patients context-specific microbiome models highlighted patient-specific variability in metabolite flux. Metabolites such as uracil and L-leucine exhibited high exchange rates in samples from patients F01, F02, and E01, while succinic acid and glycine presented consistent exchanges across multiple samples, underscoring their central roles in the metabolic network across multiple patients. Variable exchange rates of xanthosine, L-glutamic acid, and acetic acid were observed, with notable trends in samples H25363 and H25364. These findings highlight the dynamic and individualized nature of microbial metabolism, shaped by patient-specific microbiomes and their interaction with host environments.

The log-normalized rate of metabolite exchange across our entire cohort of context-specific microbiome community models was investigated, with the goal of identifying trends in metabolite exchange that are unique in specific patients or shared across the entire cohort. Our analysis highlighted significant variability in the exchange of numerous metabolites, with some metabolites showing a high metabolic flux in certain patient samples (**Figure 6B**). For instance, metabolites like uracil and L-leucine exhibited particularly high exchange rates in samples from patients F01, F02, and E01, suggesting that utilization of metabolic pathways to use these compounds are especially enriched in these patients. Succinic acid and glycine, key intermediates in microbial metabolism, were consistently exchanged at high rates across several samples; based on previously presented results, succinic acid appears to play a centralized role in the metabolic network of the UTI microbiomes through cross-feeding, linkening to it’s role as a pro-inflammtory mediator (Anthamatten et al. 2024). Other metabolites, such as xanthosine, L-glutamic acid, and acetic acid, showed variable exchange rates, with specific samples like H25363 and H25364 showing high consumption or production. These patterns suggest that certain uromicrobiomes may be more specialized than others in the production or consumption of specific metabolites, potentially influenced by the host’s metabolic environment or the microbial species present. Simulated results derived from context-specific microbiome community cross feeding for each patient can be found in **Supplementary Table S5**. The observed differences in metabolite exchange across patients highlight the dynamic and individualized nature of microbial metabolism.

## IV. Discussion

The outdated belief that the urinary tract is sterile has been recently refuted, acknowledging that a unique microbiome exists within the urinary tract, even in the absence of infections (Ackerman & Toby, 2019; Pohl et al. 2020). Our study leverages metatranscriptomics to delve into the urinary microbiome of women experiencing UTIs, revealing a complex network of microbial compositions, functions, and interactions. Contrary to the notion of UTIs being caused by single pathogens, the findings presented in this manuscript and observed by others suggest a communal and synergistic bacterial interplay (de Vos et al. 2017; Zandbergen et al. 2021). Through the integration of metatranscriptomics to reconstruct GEMs and further metabolic constraints through the Human Urine Metabolome database, personalized microbiome models for patients were reconstructed (Bouatra et al. 2013). This approach enhances our comprehension of the metabolic dynamics within the UTI microbiome, highlighting the microbial diversity and complexity in the urinary tracts of patients with UTIs. Recognizing these intricate microbial interactions is pivotal for devising more effective, patient-specific treatments.

The diversity in the uromicrobiome among UTI patients highlights the need for personalized treatment approaches, challenging the one-size-fits-all nature of standard therapies. This diversity, encompassing a range of uropathogenic, oral-associated, gut-associated, and probiotic genera, suggest that UTIs may emerge from complex microbial interactions rather than from a single pathogen. Such variability can influence immune responses, infection susceptibility, and treatment outcomes, with factors like genetics, diet, age, and antibiotic use playing significant roles in disease onset (Zheng et al. 2020; Schembri et al. 2022). The *Lactobacillus* genus, in particular, may provide protective benefits by maintaining urinary tract health and preventing pathogen colonization (Singhal et al. 2020). The Shannon index was useful to measure patient-specific diversity within patient uromicrobiomes, an approach that can be readily applicable to other microbiomes across the body (Huttenhower et al. 2012; Saheb et al. 2022). However, traditional diversity metrics may overlook the impact of low-abundance species within the uromicrobiome on community dynamics and disease progression. These minor constituents, crucial for understanding urinary health and disease, necessitate more nuanced analytical methods to fully appreciate their roles (Feehily et al. 2020; Pust et al. 2022). Our findings also point to the presence of gut-associated microbes like *E. coli and S. sonnei* in the urinary tract, underscoring a link between gastrointestinal and urogenital microbiomes in UTI development; this observation supports the hypothesis of microbial migration between these sites, though it necessitates cautious interpretation without comparative fecal sample analysis (Paalanne et al. 2018; Magruder et al. 2019; Schembri et al. 2022; Worby et al. 2022). Furthermore, we encountered minor discrepancies in sequencing data from Illumina MiSeq and NovaSeq platforms, due to varying sequencing depths and biases, which highlights a challenge in cross-platform microbial analysis (Singer et al. 2019).

The analysis of the top 10 expressed genes for UPEC UTI89 across multiple patient samples highlighted both conserved and variable transcriptional patterns. The consistent expression of *ssrA,* which is conserved across most prokaryotes, underscores its potential importance and a regulatory factor in UPEC’s cellular processes. Similarly, the frequent detection of *cspA*, and *ssrS* indicates a conserved core stress response and transcriptional regulatory role within UPEC UTI89 (Burenina et al. 2022; Cardoza et al. 2022). These findings emphasize the homeostatic tendencies of specific genes, which may be critical for baseline survival and functionality in host environments. However, the variability in the expression of other genes suggests that UPEC adapts its transcriptional profile to accommodate distinct host-specific pressures (Thänert et al. 2022). The high expression observed in UPEC UTI89 locus tags lacking assigned gene names suggests these regions may hold significant biological and potential clinical roles. Characterization of these loci through approaches such as the *E. coli* ‘y-ome’, which may be useful uncover novel molecular functions, mechanisms, or biomarkers, particularly in contexts like immune responses or disease progression (Ghatak et al. 2019 ; Moore et al. 2024). This consistent expression pattern is of potential importance for UTIs and warrants further targeted investigation to uncover their roles and improve genome annotations. UPEC strains display diverse virulence tactics, with variable expression of virulence genes among UTI patients, highlighting their adaptability and posing challenges for infection management due to different pathogenic mechanisms (Hagan et al. 2010; Frick-Cheng et al. 2020). Specific cell adherence genes, *fimA* and *fimI*, show high expression in certain samples, suggesting these strains have enhanced capabilities to invade host cells and contribute to the persistence of infections (Schüroff et al. 2021). Genes like *chuY* and *chuS*, involved in heme utilization, and iroN and fepA, related to iron acquisition, underline UPEC’s strategies to secure essential nutrients in the iron-scarce urinary tract, crucial for their virulence and survival (Mey et al. 2021; Edison et al. 2023; Zhou et al. 2023). This multifaceted virulence approach, combining adherence, invasion, and nutritional strategies, may contribute to the severity and treatment resistance of infections (Ambite et al. 2021). However, the influence of host factors on gene expression and the functional activity of expressed virulence factors remain important considerations in understanding UPEC pathogenicity. Context-specific metabolic reconstructions revealed substantial variability in UPEC UTI89’s metabolic networks across patient samples. Strains exhibited a broad range of reactions and metabolites, with some presenting expansive metabolic networks exceeding 2,000 reactions and 1,800 metabolites, while others demonstrated more constrained networks. This diversity suggests that even strains with conserved gene expression profiles employ distinct metabolic strategies (Lempp et al. 2019; Islam et al. 2022; Morrison et al. 2024). These differences likely reflect adaptive mechanisms enabling UPEC to optimize its metabolic functions to thrive in unique host environments, further underscoring the pathogen’s capacity for metabolic plasticity. However, this observation may be limited by the fact that RNA extraction success may simply differ between patient microbiomes and could be a technical limitation. Metabolic subsystem activities derived from reconstructed UPEC UTI89 models exhibited patient-specific variations, even though overall gene expression patterns were largely conserved. Subsystems such as ‘Arginine and proline metabolism,’ ‘Drug metabolism,’ ‘Glycolysis/gluconeogenesis’, and ‘Pentose phosphate pathway’ displayed variable activity across strains, with notable enrichment in strains from certain patients (e.g., A02 and C02) but limited activity in others (e.g., D01); these metabolic factors likely enhance UPEC UTI89s fitness and virulence in host environments (Alteri et al. 2009; Hibbing et al. 2020; Puebla-Barragan et al. 2020; Islam et al. 2022). Collectively, these findings underscore the need for metabolic-based approaches to complement gene expression studies, as they provide critical insights into the strain-specific adaptability of UPEC UTI89 and its implications for personalized treatment strategies.

The comparison of context-specific and non-context-specific microbiome community models revealed significant metabolic differences across pathways, highlighting the value of context-specific approaches in microbial metabolic modeling (Richelle et al., 2019b; Joshi et al., 2020; Vieira et al., 2022). Subsystems such as ‘Biosynthesis of siderophore group nonribosomal peptides,’ ‘Fatty acid synthesis,’ and ‘Glycolysis/gluconeogenesis’ exhibited substantial changes in activity under context-specific conditions, emphasizing their utilizing environmental metabolites to incite virulence mechanisms (Banerjee et al. 2020; Hong et al. 2022; Islam et al. 2022). The reduced variability and total metabolic flux observed in context-specific models, as shown by PCA and flux analyses, reflect a more biologically relevant focus on essential metabolic pathways (Joshi et al. 2020). These findings can be applied in future studies to tailor metabolic models to environmental contexts, as it was found to improve the resolution and interpretability of microbiome-wide metabolic activity. Variability in metabolic activity across distinct *Lactobacillus* community types highlights the role of microbial composition in shaping microbiome metabolism. The highest metabolic activity was observed in the *Lactobacillus Absent* group, followed by the *Lactobacillus Single* and *Lactobacillus Diverse* groups, under both context-specific and non-context-specific conditions. Statistical analyses found significant differences between *Lactobacillus Diverse* and the other two groups, suggesting that higher *Lactobacillus* diversity is associated with lower metabolic activity. Additionally, non-context-specific models consistently presented higher activity than their context-specific counterparts, further demonstrating the need for context-specific approaches to accurately represent biologically relevant metabolic activity. These results have identified the potential of a more complex relationship between the diversity of Lactobacillus in UTIs and their metabolic function in the context of uropathogens; these observations require further investigation (Song et al. 2022; Neugent et al. 2024). Analysis of log fold changes in metabolic subsystem activity between context-specific and non-context-specific conditions identified substantial variability in specific pathways, highlighting the influence of constraining microbial metabolism with gene expression. Notably, O-glycan degradation and bile acid metabolism showed the highest variability in context-specific models, particularly in samples F02, H25362, and H25363. These elevated activities suggest that these pathways are likely important underlying factors for pathogens to adapt within specific host environments that may be related to dysfunction in other organ systems, which can support pathogen binding to host cell surfaces(Sato et al. 2020; Vitko et al. 2022; Mediati et al. 2024). Conversely, subsystems like sphingolipid metabolism exhibited minimal activity in these samples, but higher in others, indicating potential niche-specific metabolic constraints related to biofilm formation in specific uropathogenic communities (Sarshar et al. 2020). These results further suggest the expanded benefits of integrating context-specific data to identify strain-specific metabolic pathways that drive microbial adaptability and pathogenicity in diverse host ecosystems.

The analysis of positive and negative flux reactions within microbial communities insights into the metabolic strategies employed by urogenital microbiomes. Positive flux reactions, such as Ferredoxin production, Cholate Transport via Bicarbonate Antiport, and Glycerol-3-phosphate acyltransferase activity, suggesting the regulation of metabolic pathways to acquire energy sources from host environments to optimize microbial survival and community stability (Vitko et al. 2022; Mediati et al. 2024; Zahn et al. 2024). In contrast, high consumption fluxes in reactions such as Hypoxanthine Exchange, Cholanate Exchange, and 5-dehydro-D-gluconate reductase highlight the reliance of these microbial communities on nutrient uptake for the synthesis pathogen lipopolysaccharide, which would rely on the scavenging of purines and the metabolism of sugars (Alteri et al. 2009; Eberly et al. 2020; Lupo et al. 2021; Heidari et al. 2024; Ljubetic et al. 2024). These findings present new insight into the microbial community’s ability to dynamically regulate its metabolic processes to optimize resource acquisition and growth to efficiently adapt within the host environment. The production and uptake of specific metabolites also revealed the nuanced metabolic interplay within urogenital microbial communities. Fermentation byproducts, including Carbon dioxide, Ammonium, Ethanol, and L-Lactate, are the most enriched metabolic outputs, as are their expected roles in anaerobic metabolism of microbial communities. The significant production of D-Alanine emphasizes its importance in the biosynthesis of cell wall components, crucial for microbial structural integrity and drug resistance in known pathogens (Jiang et al. 2021). Additionally, the high levels of Deoxyadenosine diphosphate and Guanosine production suggest an elevated demand for purine biosynthesis, integral to nucleic acid metabolism and growth (Ljubetic et al. 2023; Tan et al. 2024). On the consumption side, Hypoxanthine uptake highlights the reliance on purine salvage pathways, while the significant consumption of 7-Oxocholate and L-Idonate reflects the microbial community’s interaction with host-derived metabolites and carbohydrate metabolism (Sertbas et al. 2020; Tan et al. 2024). These finely tuned metabolic processes illustrate the complex metabolic networks within the microbial community, where production and uptake are optimized for survival and adaptation within the host environment. This regulatory balance ensures the functionality and resilience of the core urogenital microbiome.

The analysis of metabolic cross-feeding interactions within patient-specific microbial communities highlights the exchange of metabolites between community members, revealing key roles for both probiotic and pathogenic species. *Lactobacillus* species, including *L. rhamnosus* and *L. reuteri*, were identified as major producers of Alpha-lactose and SCFAs, which play maintenance roles as metabolites to balance pH in both gut and urogenital microenvironments. The production of acetate and its subsequent uptake by other uropathogens illustrates potential cross-feeding dynamics, where metabolites released by probiotics may inadvertently support pathogenic species’ adaptation and growth in the urinary tract (Elhenawy et al. 2019; Zangara et al. 2023). We also observed the unique production of ornithine by UTI89 and the consumption by the remaining members of the uromicrobiome, a metabolite which has been previously observed to provide a fitness advantage to pathogens (Durand et al. 2009; Hibbing et al. 2020). Additional findings, such as *L. amylolyticus* producing L-threonine, hydrogen, and succinic acid, underscore the potential contributions of probiotic strains to the uropathogen community metabolism (Brauer et al. 2022; Morrison et al. 2024). Interestingly, both the probiotic *L. iners* and the gut-associated *B. intestinihominis*, produce and consume metabolites like taurine and glycine, further emphasizing the complexity of cross-feeding interactions (Neugent et al. 2024). These results are important to the understanding of the individual contributing members of interspecies uropathogenic communities and their use of metabolite exchange to maintain stability across host environments. The variability in metabolite exchange rates across patient-specific microbiome models highlights the highly individualized nature of patient-specific microbial metabolism. High exchange rates of metabolites such as uracil and L-leucine in patients F01, F02, and E01 suggest that uropathogens utilize the enrichment of these metabolic pathways in specific individuals to regulate the metabolic factors to salvage pyrimidines and uptake amino-acids from host-specific environments to incite anti-persister characteristics (Wang et al. 2023; Urs et al. 2024). Succinic acid emerged as a central metabolite, consistently exhibiting high exchange rates across several samples, indicating its central role in the metabolic networks of uromicrobiomes and potentially associated with being a pro-inflammatory mediator (Anthamatten et al. 2024; Morrison et al. 2024). These observations reflect the dynamic exchanges between not only microbial communities, but also their metabolic environments which are influenced by individual host conditions. The observed differences in metabolite exchange, driven by context-specific microbial activity, reinforce the importance of personalized approaches in understanding microbial community dynamics and their implications for host health.

## V. Conclusion

Our data-driven modeling of the UTI microbiome through metatranscriptomics has provided valuable metabolic insights into metabolic pathways and cross-feeding interactions relevant to UTIs. This paves the way for the design of specific intervention strategies that interrupt those interactions e.g. via dietary supplements to suppress infection (Yang et al. 2020). However, it’s crucial to recognize the limitations of our current model. The unique phenotypes of different strains can be influenced by individual factors and micro-environmental conditions—such as diet, pre- or co-existing health conditions, local microbiota, pH, and nutrient presence—which our current framework does not entirely encapsulate. As we delve deeper into defining the functional uromicrobiome, future models will factor in patient-specific dietary and metabolic information, refining future simulations and revolutionizing how clinicians perceive and manage UTIs. Our study highlights the capability of metatranscriptomics in revealing the dynamic interplay of the uromicrobiome and suggests a roadmap for future research. As we decipher these intricate relationships between the host and its microbiome in UTIs, the vision of tailoring therapeutic strategies based on individual microbiome compositions emerges, heralding a new era of enhanced care for UTI patients.

## Acknowledgments

The authors are thankful to the Medical Systems Biology team for useful discussions and support in this project. The graphical abstract was created using Biorender. The authors declare that they have no conflict of interest.

## DATA/CODE AVAILABILITY

All in-house generated sequences were deposited in the NCBI-Sequence Read Archive database (SRP498164). The accession number of the deposited reads is provided in **Supplementary Table S1**. Code/data/software required to replicate the figures and findings in this manuscript are all available on GitHub (https://github.com/PoeticPremium6/UTI-Microbiome-Model).

## References

Ackerman, A. Lenore, and Toby C. Chai. “The bladder is not sterile: an update on the urinary microbiome.” Current bladder dysfunction reports 14 (2019): 331–341.

Alteri, Christopher J., Sara N. Smith, and Harry LT Mobley. “Fitness of Escherichia coli during urinary tract infection requires gluconeogenesis and the TCA cycle.” PLoS pathogens 5, no. 5 (2009): e1000448.

Ambite, Ines, Daniel Butler, Murphy Lam Yim Wan, Therese Rosenblad, Thi Hien Tran, Sing Ming Chao, and Catharina Svanborg. “Molecular determinants of disease severity in urinary tract infection.” Nature Reviews Urology 18, no. 8 (2021): 468–486.

Anthamatten, Laura, Philipp Rogalla von Bieberstein, Carmen Menzi, Janina N. Zünd, Christophe Lacroix, Tomas de Wouters, and Gabriel E. Leventhal. “Stratification of human gut microbiomes by succinotype is associated with inflammatory bowel disease status.” Microbiome 12, no. 1 (2024): 186.

Baker, Joseph L., Tobias Dahlberg, Esther Bullitt, and Magnus Andersson. “Impact of an alpha helix and a cysteine–cysteine disulfide bond on the resistance of bacterial adhesion pili to stress.” Proceedings of the National Academy of Sciences 118, no. 21 (2021): e2023595118.

Banerjee, Rajdeep, Erin Weisenhorn, Kevin J. Schwartz, Kevin S. Myers, Jeremy D. Glasner, Nicole T. Perna, Joshua J. Coon, Rodney A. Welch, and Patricia J. Kiley. “Tailoring a global iron regulon to a uropathogen.” MBio 11, no. 2 (2020): 10–1128.

Bauer, Eugen, Johannes Zimmermann, Federico Baldini, Ines Thiele, and Christoph Kaleta. “BacArena: Individual-based metabolic modeling of heterogeneous microbes in complex communities.” PLoS computational biology 13, no. 5 (2017): e1005544.

Bouatra, Souhaila, Farid Aziat, Rupasri Mandal, An Chi Guo, Michael R. Wilson, Craig Knox, Trent C. Bjorndahl et al. “The human urine metabolome.” PloS one 8, no. 9 (2013): e73076.

Brauer, Aimee L., Brian S. Learman, Steven M. Taddei, Namrata Deka, Benjamin C. Hunt, and Chelsie E. Armbruster. “Preferential catabolism of l-vs d-serine by Proteus mirabilis contributes to pathogenesis and catheter-associated urinary tract infection.” Molecular microbiology 118, no. 3 (2022): 125–144.

Burenina, Olga Y., Daria A. Elkina, Anna Ovcharenko, Valeria A. Bannikova, M. Amri C. Schlüter, Tatiana S. Oretskaya, Roland K. Hartmann, and Elena A. Kubareva. “Involvement of E. coli 6S RNA in oxidative stress response.” International Journal of Molecular Sciences 23, no. 7 (2022): 3653.

Cardoza, Evieann, and Harinder Singh. “Involvement of CspC in response to diverse environmental stressors in Escherichia coli.” Journal of Applied Microbiology 132, no. 2 (2022): 785–801.

Danecek, Petr, James K. Bonfield, Jennifer Liddle, John Marshall, Valeriu Ohan, Martin O. Pollard, Andrew Whitwham et al. “Twelve years of SAMtools and BCFtools.” Gigascience 10, no. 2 (2021): giab008.

Deng, Zhi-Luo, Cornelia Gottschick, Sabin Bhuju, Clarissa Masur, Christoph Abels, and Irene Wagner-Döbler. “Metatranscriptome analysis of the vaginal microbiota reveals potential mechanisms for protection against metronidazole in bacterial vaginosis.” Msphere 3, no. 3 (2018): 10–1128.

de Vos, Marjon GJ, Marcin Zagorski, Alan McNally, and Tobias Bollenbach. “Interaction networks, ecological stability, and collective antibiotic tolerance in polymicrobial infections.” Proceedings of the National Academy of Sciences 114, no. 40 (2017): 10666–10671.

Durand, Jérôme MB, and Glenn R. Bjork. “Metabolic control through ornithine and uracil of epithelial cell invasion by Shigella flexneri.” Microbiology 155, no. 8 (2009): 2498–2508.

Eberly, Allison R., Connor J. Beebout, Ching Man Carmen Tong, Gerald T. Van Horn, Hamilton D. Green, Madison J. Fitzgerald, Shuvro De et al. ”Defining a molecular signature for uropathogenic versus urocolonizing Escherichia coli: the status of the field and new clinical opportunities.” Journal of molecular biology 432, no. 4 (2020): 786–804.

Edgar, Robert. Usearch. Lawrence Berkeley National Lab.(LBNL), Berkeley, CA (United States), 2010.

Edison, Lekshmi K., Indira T. Kudva, and Subhashinie Kariyawasam. “Comparative Transcriptome Analysis of Shiga Toxin-Producing Escherichia coli O157: H7 on Bovine Rectoanal Junction Cells and Human Colonic Epithelial Cells during Initial Adherence.” Microorganisms 11, no. 10 (2023): 2562.

Elhenawy, Wael, Caressa N. Tsai, and Brian K. Coombes. “Host-specific adaptive diversification of Crohn’s disease-associated adherent-invasive Escherichia coli.” Cell Host & Microbe 25, no. 2 (2019): 301–312.

Feehily, Conor, David Crosby, Calum J. Walsh, Elaine M. Lawton, Shane Higgins, Fionnuala M. McAuliffe, and Paul D. Cotter. “Shotgun sequencing of the vaginal microbiome reveals both a species and functional potential signature of preterm birth.” NPJ biofilms and microbiomes 6, no. 1 (2020): 50.

Flores-Mireles, Ana L., Jennifer N. Walker, Michael Caparon, and Scott J. Hultgren. “Urinary tract infections: epidemiology, mechanisms of infection and treatment options.” Nature reviews microbiology 13, no. 5 (2015): 269–284.

Frick-Cheng, Arwen E., Anna Sintsova, Sara N. Smith, Michael Krauthammer, Kathryn A. Eaton, and Harry LT Mobley. “The gene expression profile of uropathogenic Escherichia coli in women with uncomplicated urinary tract infections is recapitulated in the mouse model.” MBio 11, no. 4 (2020): 10–1128.

Fu, Limin, Beifang Niu, Zhengwei Zhu, Sitao Wu, and Weizhong Li. “CD-HIT: accelerated for clustering the next-generation sequencing data.” Bioinformatics 28, no. 23 (2012): 3150–3152.

George, Sheridan D., Olivia T. Van Gerwen, Chaoling Dong, Lúcia GV Sousa, Nuno Cerca, Jacob H. Elnaggar, Christopher M. Taylor, and Christina A. Muzny. “The Role of Prevotella Species in Female Genital Tract Infections.” Pathogens 13, no. 5 (2024): 364.

Ghatak, Sankha, Zachary A. King, Anand Sastry, and Bernhard O. Palsson. “The y-ome defines the 35% of Escherichia coli genes that lack experimental evidence of function.” Nucleic acids research 47, no. 5 (2019): 2446–2454.

Gopalakrishnan, Saratram, Chintan J. Joshi, Miguel Á. Valderrama-Gómez, Elcin Icten, Pablo Rolandi, William Johnson, Cleo Kontoravdi, and Nathan E. Lewis. “Guidelines for extracting biologically relevant context-specific metabolic models using gene expression data.” Metabolic Engineering 75 (2023): 181–191.

Graspeuntner, Simon, Nathalie Loeper, Sven Künzel, John F. Baines, and Jan Rupp. “Selection of validated hypervariable regions is crucial in 16S-based microbiota studies of the female genital tract.” Scientific reports 8, no. 1 (2018): 9678.

Graspeuntner, Simon, Mariia Lupatsii, Vera van Zandbergen, Marie-Theres Dammann, Julia Pagel, Duc Ninh Nguyen, Alexander Humberg et al. “Infants< 90 days of age with late-onset sepsis display disturbances of the microbiome-immunity interplay.” Infection (2024): 1–14.

Hagan, Erin C., Amanda L. Lloyd, David A. Rasko, Gary J. Faerber, and Harry LT Mobley. “Escherichia coli global gene expression in urine from women with urinary tract infection.” PLoS pathogens 6, no. 11 (2010): e1001187.

Heidari, Aida, Mohammad Hassan Emami, Fatemeh Maghool, Samane Mohammadzadeh, Parisa Kadkhodaei Elyaderani, Tahereh Safari, Alireza Fahim, and Razie Kamali Dolatabadi. “Molecular epidemiology, antibiotic resistance profile and frequency of integron 1 and 2 in adherent-invasive Escherichia coli isolates of colorectal cancer patients.” Frontiers in Microbiology 15 (2024): 1366719.

Heirendt, Laurent, Sylvain Arreckx, Thomas Pfau, Sebastián N. Mendoza, Anne Richelle, Almut Heinken, Hulda S. Haraldsdóttir et al. “Creation and analysis of biochemical constraint-based models using the COBRA Toolbox v. 3.0.” Nature protocols 14, no. 3 (2019): 639–702.

Heinken, Almut, Johannes Hertel, Geeta Acharya, Dmitry A. Ravcheev, Malgorzata Nyga, Onyedika Emmanuel Okpala, Marcus Hogan et al. “Genome-scale metabolic reconstruction of 7,302 human microorganisms for personalized medicine.” Nature Biotechnology (2023): 1–12.

Hibbing, Michael E., Karen W. Dodson, Vasilios Kalas, Swaine L. Chen, and Scott J. Hultgren. “Adaptation of arginine synthesis among uropathogenic branches of the Escherichia coli phylogeny reveals adjustment to the urinary tract habitat.” MBio 11, no. 5 (2020): 10–1128.

Hong, Yaoqin, Jilong Qin, Anthony D. Verderosa, Sophia Hawas, Bing Zhang, Mark AT Blaskovich, John E. Cronan Jr, and Makrina Totsika. “Loss of β-ketoacyl acyl carrier protein synthase III activity restores multidrug-resistant Escherichia coli sensitivity to previously ineffective antibiotics.” Msphere 7, no. 3 (2022): e00117–22.

Hooton, Thomas M. “Uncomplicated urinary tract infection.” New England Journal of Medicine 366, no. 11 (2012): 1028–1037.

Huttenhower, Curtis, Dirk Gevers, Rob Knight, Sahar Abubucker, Jonathan H. Badger, Asif T. Chinwalla, Heather H. Creasy et al. “Structure, function and diversity of the healthy human microbiome.” Nature 486, no. 7402 (2012): 207–214.

Islam, Md Jahirul, Kamal Bagale, Preeti P. John, Zachary Kurtz, and Ritwij Kulkarni. “Glycosuria alters uropathogenic Escherichia coli global gene expression and virulence.” Msphere 7, no. 3 (2022): e00004–22.

Jiang, Qingsong, Xiaoya He, Yusen Shui, Xiaoying Lyu, Liang Wang, Laijun Xu, Zhu Chen et al. “d-Alanine metabolic pathway, a potential target for antibacterial drug designing in Enterococcus faecalis.” Microbial Pathogenesis 158 (2021): 105078.

Joshi, Chintan J., Song-Min Schinn, Anne Richelle, Isaac Shamie, Eyleen J. O’Rourke, and Nathan E. Lewis. “StanDep: Capturing transcriptomic variability improves context-specific metabolic models.” PLoS computational biology 16, no. 5 (2020): e1007764.

Kalvari, Ioanna, Eric P. Nawrocki, Nancy Ontiveros-Palacios, Joanna Argasinska, Kevin Lamkiewicz, Manja Marz, Sam Griffiths-Jones et al. “Rfam 14: expanded coverage of metagenomic, viral and microRNA families.” Nucleic Acids Research 49, no. D1 (2021): D192–D200.

Kim, Daehwan, Joseph M. Paggi, Chanhee Park, Christopher Bennett, and Steven L. Salzberg. “Graph-based genome alignment and genotyping with HISAT2 and HISAT-genotype.” Nature biotechnology 37, no. 8 (2019): 907–915.

Kopylova, Evguenia, Laurent Noé, and Hélène Touzet. “SortMeRNA: fast and accurate filtering of ribosomal RNAs in metatranscriptomic data.” Bioinformatics 28, no. 24 (2012): 3211–3217.

Langmead, Ben, and Steven L. Salzberg. “Fast gapped-read alignment with Bowtie 2.” Nature methods 9, no. 4 (2012): 357–359.

Lempp, Martin, Niklas Farke, Michelle Kuntz, Sven Andreas Freibert, Roland Lill, and Hannes Link. “Systematic identification of metabolites controlling gene expression in E. coli.” Nature communications 10, no. 1 (2019): 4463.

Liao, Yang, Gordon K. Smyth, and Wei Shi. “featureCounts: an efficient general purpose program for assigning sequence reads to genomic features.” Bioinformatics 30, no. 7 (2014): 923–930.

Liu, Bo, Dandan Zheng, Siyu Zhou, Lihong Chen, and Jian Yang. “VFDB 2022: a general classification scheme for bacterial virulence factors.” Nucleic acids research 50, no. D1 (2022): D912–D917.

Ljubetic, Bernardita M., Ashu Mohammad, Butool Durrani, and Amy D. Dobberfuhl. “Pathophysiologic Insights into the Transition from Asymptomatic Bacteriuria to Urinary Tract Infection.” Current Urology Reports 24, no. 11 (2023): 533–540.

Lupo, Federico, Molly A. Ingersoll, and Miguel A. Pineda. “The glycobiology of uropathogenic E. coli infection: the sweet and bitter role of sugars in urinary tract immunity.” Immunology 164, no. 1 (2021): 3–14.

Lussu, Milena, Tania Camboni, Cristina Piras, Corrado Serra, Francesco Del Carratore, Julian Griffin, Luigi Atzori, and Aldo Manzin. “1 H NMR spectroscopy-based metabolomics analysis for the diagnosis of symptomatic E. coli-associated urinary tract infection (UTI).” BMC microbiology 17 (2017): 1–8.

Lüth, Theresa, Simon Graspeuntner, Kay Neumann, Laura Kirchhoff, Antonia Masuch, Susen Schaake, Mariia Lupatsii et al. “Improving analysis of the vaginal microbiota of women undergoing assisted reproduction using nanopore sequencing.” Journal of Assisted Reproduction and Genetics 39, no. 11 (2022): 2659–2667.

Magruder, Matthew, Adam N. Sholi, Catherine Gong, Lisa Zhang, Emmanuel Edusei, Jennifer Huang, Shady Albakry et al. “Gut uropathogen abundance is a risk factor for development of bacteriuria and urinary tract infection.” Nature communications 10, no. 1 (2019): 5521.

Marinos, Georgios, Christoph Kaleta, and Silvio Waschina. “Defining the nutritional input for genome-scale metabolic models: A roadmap.” PloS one 15, no. 8 (2020): e0236890.

Martin, Marcel. “Cutadapt removes adapter sequences from high-throughput sequencing reads.” *EMBnet*. journal 17, no. 1 (2011): 10–12.

McMurdie, Paul J., and Susan Holmes. “phyloseq: an R package for reproducible interactive analysis and graphics of microbiome census data.” PloS one 8, no. 4 (2013): e61217.

Mediati, Daniel G., Tamika A. Blair, Ariana Costas, Leigh G. Monahan, Bill Söderström, Ian G. Charles, and Iain G. Duggin. “Genetic requirements for uropathogenic E. coli proliferation in the bladder cell infection cycle.” Msystems 9, no. 10 (2024): e00387–24.

Mediavilla, José R., Amee Patrawalla, Liang Chen, Kalyan D. Chavda, Barun Mathema, Christopher Vinnard, Lisa L. Dever, and Barry N. Kreiswirth. “Colistin-and carbapenem-resistant Escherichia coli harboring mcr-1 and bla NDM-5, causing a complicated urinary tract infection in a patient from the United States.” MBio 7, no. 4 (2016): 10–1128.

Mey, Alexandra R., Camilo Gómez-Garzón, and Shelley M. Payne. “Iron transport and metabolism in Escherichia, Shigella, and Salmonella.” EcoSal plus 9, no. 2 (2021): eESP–0034.

Miller, Sophie J., Lucy Carpenter, Steven L. Taylor, Steve L. Wesselingh, Jocelyn M. Choo, Andrew P. Shoubridge, Lito E. Papanicolas, Geraint B. Rogers, and GRACE Investigator Group Flynn Erin 4 Gordon David 3 4 Lynn David J. 2 5 Whitehead Craig 6 7 Leong Lex EX 2 Crotty Maria 6 7 Inacio Maria 7 8. “Intestinal microbiology and urinary tract infection associated risk in long-term aged care residents.” Communications Medicine 4, no. 1 (2024): 164.

Morrison, Josiah J., Ellen K. Madden, Daniel A. Banas, Eric C. DiBiasio, Mads Hansen, Karen A. Krogfelt, David C. Rowley, Paul S. Cohen, and Jodi L. Camberg. “Metabolic flux regulates growth transitions and antibiotic tolerance in uropathogenic Escherichia coli.” Journal of Bacteriology (2024): e00162–24.

Moore, Lisa R., Ron Caspi, Dana Boyd, Mehmet Berkmen, Amanda Mackie, Suzanne Paley, and Peter D. Karp. “Revisiting the y-ome of Escherichia coli.” Nucleic Acids Research 52, no. 20 (2024): 12201–12207.

Neugent, Michael L., Neha V. Hulyalkar, Kevin C. Lutz, Qiwei Li, Philippe E. Zimmern, Vladimir Shulaev, and Nicole J. De Nisco. “MP69-02 THE IMPACT OF RECURRENT URINARY TRACT INFECTION AND UROBIOME ECOLOGY ON THE URINARY METABOLOME.” Journal of Urology 211, no. 5S (2024): e1118.

Noronha, Alberto, Jennifer Modamio, Yohan Jarosz, Elisabeth Guerard, Nicolas Sompairac, German Preciat, Anna Dröfn Daníelsdóttir et al. “The Virtual Metabolic Human database: integrating human and gut microbiome metabolism with nutrition and disease.” Nucleic acids research 47, no. D1 (2019): D614–D624.

O’Brien, Edward J., Jonathan M. Monk, and Bernhard O. Palsson. “Using genome-scale models to predict biological capabilities.” Cell 161, no. 5 (2015): 971–987.

Ojala, Teija, Esko Kankuri, and Matti Kankainen. “Understanding human health through metatranscriptomics.” Trends in Molecular Medicine 29, no. 5 (2023): 376–389.

Paalanne, Niko, Aleksi Husso, Jarmo Salo, Oskari Pieviläinen, Mysore V. Tejesvi, Pirjo Koivusaari, Anna Maria Pirttilä et al. ”Intestinal microbiome as a risk factor for urinary tract infections in children.” European Journal of Clinical Microbiology & Infectious Diseases 37 (2018): 1881–1891.

Pan, Yongdong, Jingyi Su, Shengnan Liu, Yueyan Li, and Guofeng Xu. “Causal effects of gut microbiota on the risk of urinary tract stones: A bidirectional two-sample mendelian randomization study.” Heliyon 10, no. 4 (2024).

Pertea, Mihaela, Geo M. Pertea, Corina M. Antonescu, Tsung-Cheng Chang, Joshua T. Mendell, and Steven L. Salzberg. “StringTie enables improved reconstruction of a transcriptome from RNA-seq reads.” Nature biotechnology 33, no. 3 (2015): 290–295.

Pohl, Hans G., Suzanne L. Groah, Marcos Pérez-Losada, Inger Ljungberg, Bruce M. Sprague, Neel Chandal, Ljubica Caldovic, and Michael Hsieh. “The urine microbiome of healthy men and women differs by urine collection method.” International Neurourology Journal 24, no. 1 (2020): 41.

Pust, Marie-Madlen, and Burkhard Tümmler. “Bacterial low-abundant taxa are key determinants of a healthy airway metagenome in the early years of human life.” Computational and Structural Biotechnology Journal 20 (2022): 175–186.

Richelle, Anne, Chintan Joshi, and Nathan E. Lewis. “Assessing key decisions for transcriptomic data integration in biochemical networks.” PLoS computational biology 15, no. 7 (2019a): e1007185.

Richelle, Anne, Austin WT Chiang, Chih-Chung Kuo, and Nathan E. Lewis. “Increasing consensus of context-specific metabolic models by integrating data-inferred cell functions.” PLoS computational biology 15, no. 4 (2019b): e1006867.

Roussel, Charlène, Stéphane Chabaud, Jacob Lessard-Lord, Valentina Cattero, Félix-Antoine Pellerin, Perrine Feutry, Valérie Bochard, Stéphane Bolduc, and Yves Desjardins. “UPEC colonic-virulence and urovirulence are blunted by proanthocyanidins-rich cranberry extract microbial metabolites in a gut model and a 3D tissue-engineered urothelium.” Microbiology Spectrum 10, no. 5 (2022): e02432–21.

Russo, Filippo, Speranza Esposito, Lorella Tripodi, Savio Domenico Pandolfo, Achille Aveta, Felice Amato, Carmela Nardelli, Ciro Imbimbo, Lucio Pastore, and Giuseppe Castaldo. “Insights into Porphyromonas somerae in Bladder Cancer Patients: Urinary Detection by ddPCR.” Microorganisms 12, no. 10 (2024): 2049.

Saheb Kashaf, Sara, Diana M. Proctor, Clay Deming, Paul Saary, Martin Hölzer, Monica E. Taylor, Heidi H. Kong, Julia A. Segre, Alexandre Almeida, and Robert D. Finn. “Integrating cultivation and metagenomics for a multi-kingdom view of skin microbiome diversity and functions.” Nature microbiology 7, no. 1 (2022): 169–179.

Sato, Keiko, Genta Kakiyama, Mitsuyoshi Suzuki, Nakayuki Naritaka, Hajime Takei, Hiroaki Sato, Akihiko Kimura et al. “Changes in conjugated urinary bile acids across age groups.” Steroids 164 (2020): 108730.

Sarshar, Meysam, Payam Behzadi, Cecilia Ambrosi, Carlo Zagaglia, Anna Teresa Palamara, and Daniela Scribano. “FimH and anti-adhesive therapeutics: a disarming strategy against uropathogens.” Antibiotics 9, no. 7 (2020): 397.

Schmieder, Robert, and Robert Edwards. “Quality control and preprocessing of metagenomic datasets.” Bioinformatics 27, no. 6 (2011): 863–864.

Schembri, Mark A., Nguyen Thi Khanh Nhu, and Minh-Duy Phan. “Gut–bladder axis in recurrent UTI.” Nature Microbiology 7, no. 5 (2022): 601–602.

Schüroff, Paulo A., Fábia A. Salvador, Cecilia M. Abe, Haleluya T. Wami, Eneas Carvalho, Rodrigo T. Hernandes, Ulrich Dobrindt, Tânia AT Gomes, and Waldir P. Elias. “The aggregate-forming pili (AFP) mediates the aggregative adherence of a hybrid-pathogenic Escherichia coli (UPEC/EAEC) isolated from a urinary tract infection.” Virulence 12, no. 1 (2021): 3073–3093.

Sen, Partho, and Matej Orešič. “Integrating Omics Data in Genome-Scale Metabolic Modeling: A Methodological Perspective for Precision Medicine.” Metabolites 13, no. 7 (2023): 855.

Sertbas, Mustafa, and Kutlu O. Ulgen. “Genome-scale metabolic modeling for unraveling molecular mechanisms of high threat pathogens.” Frontiers in Cell and Developmental Biology 8 (2020): 566702.

Shields, Ryan K., Yun Zhou, Hemanth Kanakamedala, and Bin Cai. “Burden of illness in US hospitals due to carbapenem-resistant Gram-negative urinary tract infections in patients with or without bacteraemia.” BMC infectious diseases 21, no. 1 (2021): 1–12.

Singhal, Lipika, Varsha Gupta, Menal Gupta, Poonam Goel, and Jagdish Chander. “Identification and Sensitivity of Vaginal and Probiotic Lactobacillus species to Urinary Antibiotics.” Journal of Laboratory Physicians 12, no. 02 (2020): 111–114.

Singer, G. A. C., Nicole A. Fahner, J. G. Barnes, Avery McCarthy, and Mehrdad Hajibabaei. “Comprehensive biodiversity analysis via ultra-deep patterned flow cell technology: a case study of eDNA metabarcoding seawater.” Scientific reports 9, no. 1 (2019): 5991.

Song, Chang Hyun, Young Ho Kim, Manisha Naskar, Byron W. Hayes, Mathew A. Abraham, Joo Hwan Noh, Gyeongseo Suk et al. “Lactobacillus crispatus limits bladder uropathogenic E. coli infection by triggering a host type I interferon response.” Proceedings of the National Academy of Sciences 119, no. 33 (2022): e2117904119.

Tan, Casandra Ai Zhu, Kelvin Kian Long Chong, Daryl Yu Xuan Yeong, Celine Hui Min Ng, Muhammad Hafiz Ismail, Zhei Hwee Yap, Varnica Khetrapal et al. ”Purine and carbohydrate availability drive Enterococcus faecalis fitness during wound and urinary tract infections.” Mbio 15, no. 1 (2024): e02384–23.

Thänert, Robert, JooHee Choi, Kimberly A. Reske, Tiffany Hink, Anna Thänert, Meghan A. Wallace, Bin Wang et al. “Persisting uropathogenic Escherichia coli lineages show signatures of niche-specific within-host adaptation mediated by mobile genetic elements.” Cell host & microbe 30, no. 7 (2022): 1034–1047.

Thiele, Ines, Swagatika Sahoo, Almut Heinken, Johannes Hertel, Laurent Heirendt, Maike K. Aurich, and Ronan MT Fleming. “Personalized whole-body models integrate metabolism, physiology, and the gut microbiome.” Molecular systems biology 16, no. 5 (2020): e8982.

Tufon, Kukwah Anthony, Djike Puepi Yolande Fokam, Youmbi Sylvain Kouanou, and Henry Dilonga Meriki. “Case report on a swift shift in uropathogens from Shigella flexneri to Escherichia coli: a thin line between bacterial persistence and reinfection.” Annals of Clinical Microbiology and Antimicrobials 19, no. 1 (2020): 31.

Urs, Karthik, Philippe E. Zimmern, and Larry Reitzer. “Abundant urinary amino acids activate glutamine synthetase-encoding glnA by two different mechanisms in Escherichia coli.” Journal of bacteriology 206, no. 3 (2024): e00376–23.

van Teijlingen, Nienke H., Marleen Y. van Smoorenburg, Ramin Sarrami-Forooshani, Esther M. Zijlstra-Willems, John L. van Hamme, Hanneke Borgdorff, Janneke HHM van de Wijgert et al. “Prevotella timonensis Bacteria Associated With Vaginal Dysbiosis Enhance Human Immunodeficiency Virus Type 1 Susceptibility Of Vaginal CD4+ T Cells.” The Journal of Infectious Diseases (2024): jiae166.

Vieira, Vítor, Jorge Ferreira, and Miguel Rocha. “A pipeline for the reconstruction and evaluation of context-specific human metabolic models at a large-scale.” PLoS computational biology 18, no. 6 (2022): e1009294.

Vitko, Dijana, Joseph W. McQuaid, Ali Hashemi Gheinani, Kohei Hasegawa, Shannon DiMartino, Kylie H. Davis, Candace Y. Chung et al. “Urinary tract infections in children with vesicoureteral reflux are accompanied by alterations in urinary microbiota and metabolome profiles.” European Urology 81, no. 2 (2022): 151–154.

Vlassis, Nikos, Maria Pires Pacheco, and Thomas Sauter. “Fast reconstruction of compact context-specific metabolic network models.” PLoS computational biology 10, no. 1 (2014): e1003424.

Wang, Yanyan, Bing Liang, Zhengming Song, Wujun Chen, Hongxia Niu, Dongming Xing, and Ying Zhang. “High antipersister activity of a promising new quinolone drug candidate in eradicating uropathogenic Escherichia coli persisters and persistent infection in mice.” Journal of Applied Microbiology 134, no. 9 (2023): lxad193.

Wingett, Steven W., and Simon Andrews. “FastQ Screen: A tool for multi-genome mapping and quality control.” F1000Research 7 (2018).

Worby, Colin J., Henry L. Schreiber IV, Timothy J. Straub, Lucas R. van Dijk, Ryan A. Bronson, Benjamin S. Olson, Jerome S. Pinkner, et al. “Longitudinal multi-omics analyses link gut microbiome dysbiosis with recurrent urinary tract infections in women.” Nature microbiology 7, no. 5 (2022): 630–639.

Yilmaz, Pelin, Laura Wegener Parfrey, Pablo Yarza, Jan Gerken, Elmar Pruesse, Christian Quast, Timmy Schweer, Jörg Peplies, Wolfgang Ludwig, and Frank Oliver Glöckner. “The SILVA and “all-species living tree project (LTP)” taxonomic frameworks.” Nucleic acids research 42, no. D1 (2014): D643–D648.

Zahn, Leah E., Paige M. Gannon, and Lauren J. Rajakovich. “Iron-sulfur cluster-dependent enzymes and molybdenum-dependent reductases in the anaerobic metabolism of human gut microbes.” Metallomics (2024): mfae049.

Zampieri, Guido, Stefano Campanaro, Claudio Angione, and Laura Treu. “Metatranscriptomics-guided genome-scale metabolic modeling of microbial communities.” Cell Reports Methods 3, no. 1 (2023).

Zandbergen, Laurens E., Thomas Halverson, Jolanda K. Brons, Alan J. Wolfe, and Marjon GJ De Vos. “The good and the bad: ecological interaction measurements between the urinary microbiota and uropathogens.” Frontiers in Microbiology 12 (2021): 659450.

Zangara, Megan T., Lena Darwish, and Brian K. Coombes. “Characterizing the pathogenic potential of Crohn’s disease-associated adherent-invasive Escherichia coli.” EcoSal Plus 11, no. 1 (2023): eesp-0018.

Zhou, Yang, Zuying Zhou, Lin Zheng, Zipeng Gong, Yueting Li, Yang Jin, Yong Huang, and Mingyan Chi. “Urinary tract infections caused by uropathogenic Escherichia coli: mechanisms of infection and treatment options.” International journal of molecular sciences 24, no. 13 (2023): 10537.

Zheng, Danping, Timur Liwinski, and Eran Elinav. “Interaction between microbiota and immunity in health and disease.” Cell research 30, no. 6 (2020): 492–506.

Zimmermann, Johannes, Christoph Kaleta, and Silvio Waschina. “gapseq: Informed prediction of bacterial metabolic pathways and reconstruction of accurate metabolic models.” Genome biology 22, no. 1 (2021): 1–35.

