## Supplementary Figures for "Metatranscriptomics-based metabolic modeling of patient-specific urinary microbiome during infection"

**Supplementary Material**

**Supplementary Tables**

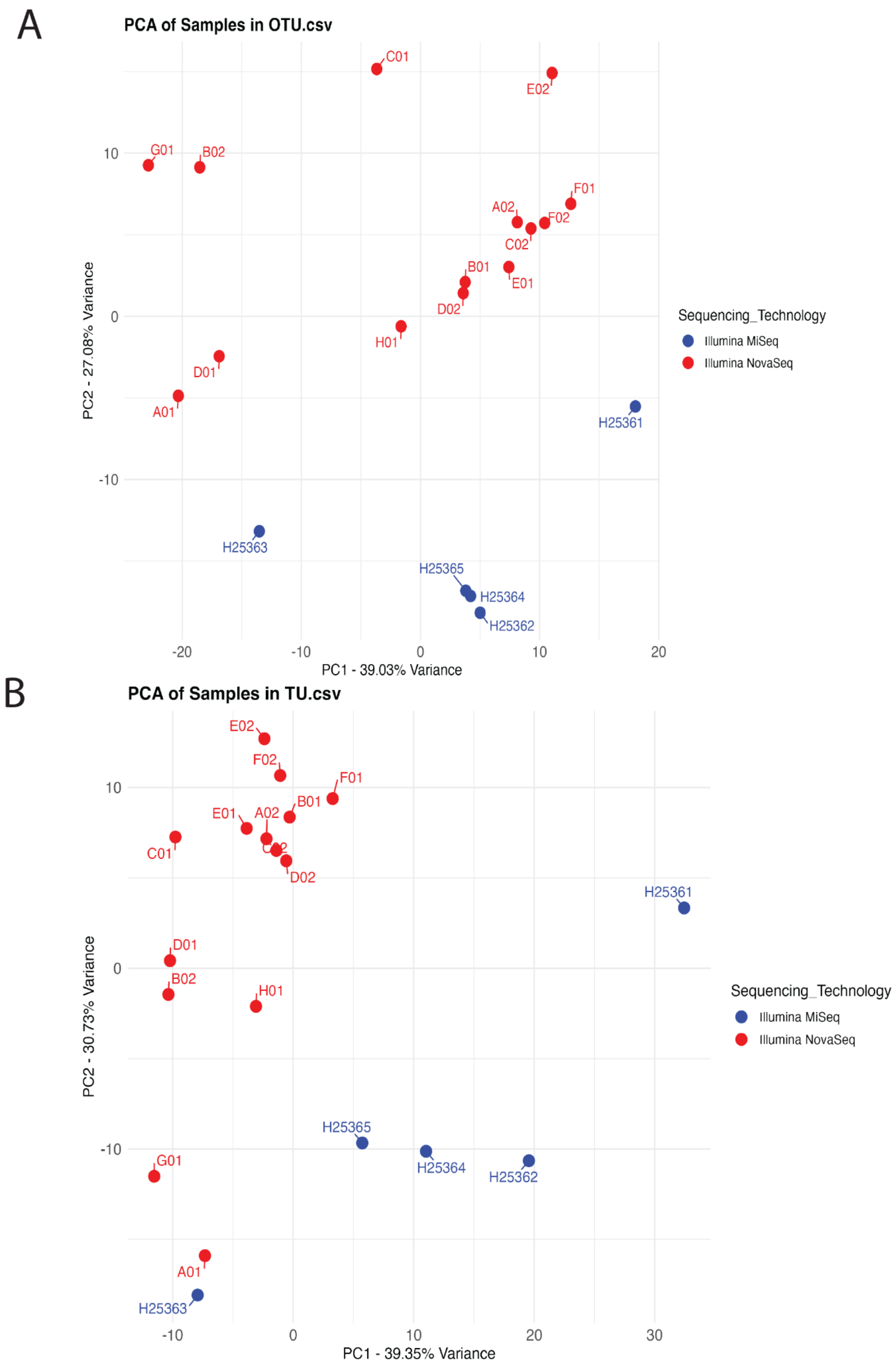

A
PCA of Samples in OTU.csv
PC1 - 39.03% Variance
PC2 - 27.08% Variance
Sequencing_Technology
Illumina MiSeq
Illumina NovaSeq
C01
E02
G01
B02
F01
A02
F02
C02
B01
E01
D02
H01
D01
A01
H25361
H25363
H25365
H25364
H25362
B
PCA of Samples in TU.csv
PC1 - 39.35% Variance
PC2 - 30.73% Variance
Sequencing_Technology
Illumina MiSeq
Illumina NovaSeq
E02
F02
F01
B01
E01
A02
C01
D02
H25361
D01
B02
H01
H25365
H25364
H25362
G01
A01
H25363

**Supplementary Figure S1**: **PCA assessment of TUs and OTUs against sequencing technology**

(A) PCA for OTUs visualizes microbial diversity, similarly coded by sequencing technology: Illumina MiSeq (blue circles) and NovaSeq (red triangles). (B) PCA plot for TUs visualzes transcription unit variance across samples by sequencing technology

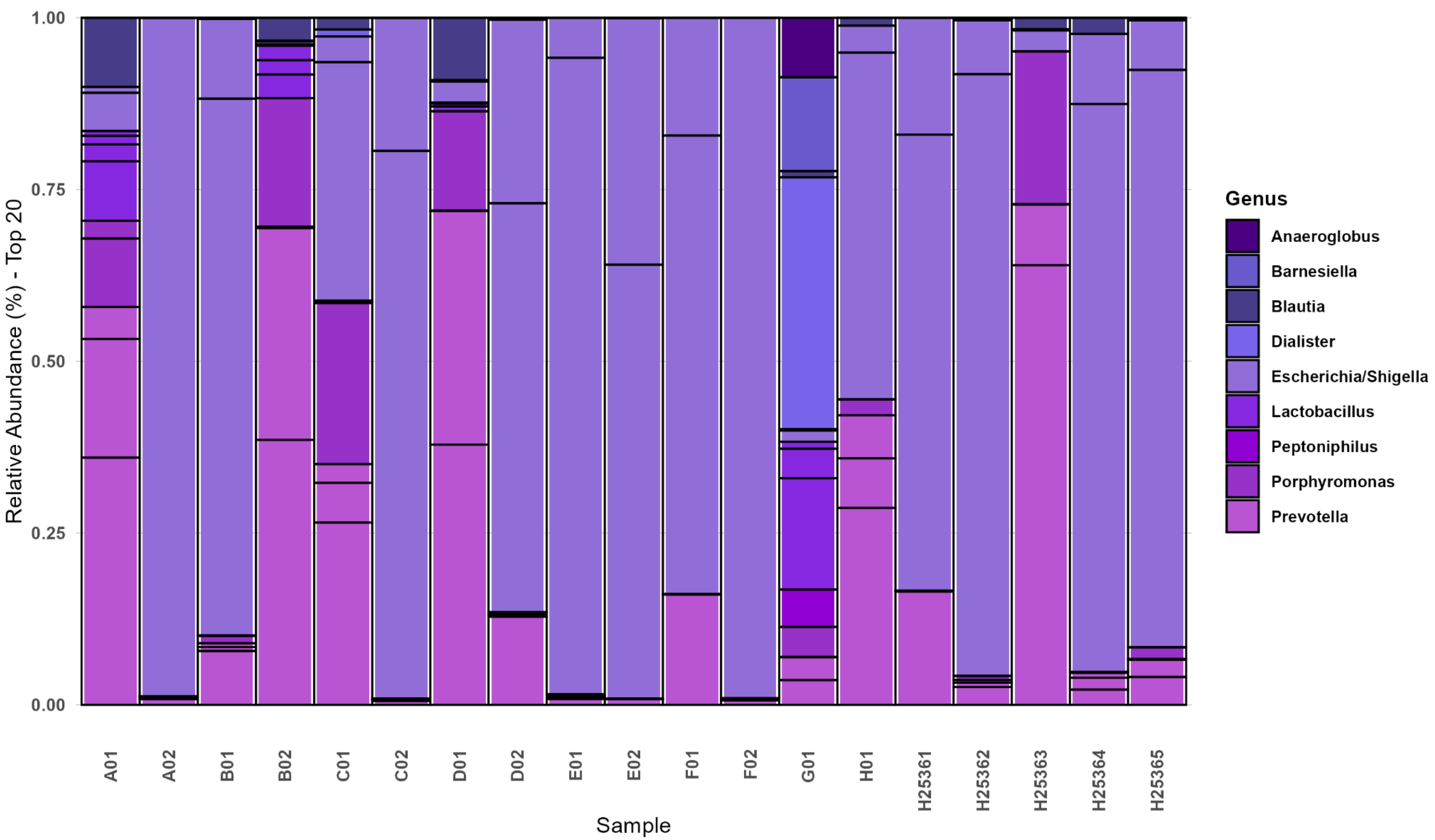

Relative Abundance (%) - Top 20
1.00
0.75
0.50
0.25
0.00
A01
A02
B01
B02
C01
C02
D01
D02
E01
E02
F01
F02
G01
H01
H25361
H25362
H25363
H25364
H25365
Sample
Genus
Anaeroglobus
Barnesiella
Blautia
Dialister
Escherichia/Shigella
Lactobacillus
Peptoniphilus
Porphyromonas
Prevotella

**Supplementary Figure S2**: **Taxonomic Abundance of Specific UTI Genera**

(A) A focused examination of six night microbial genera across our uromicrobiome consortia. This panel underscores the varying prevalence of these pathogenic genera across different patient samples, thereby highlighting the unique microbial signatures that define each UTI microbiome and their distribution among individual patients.

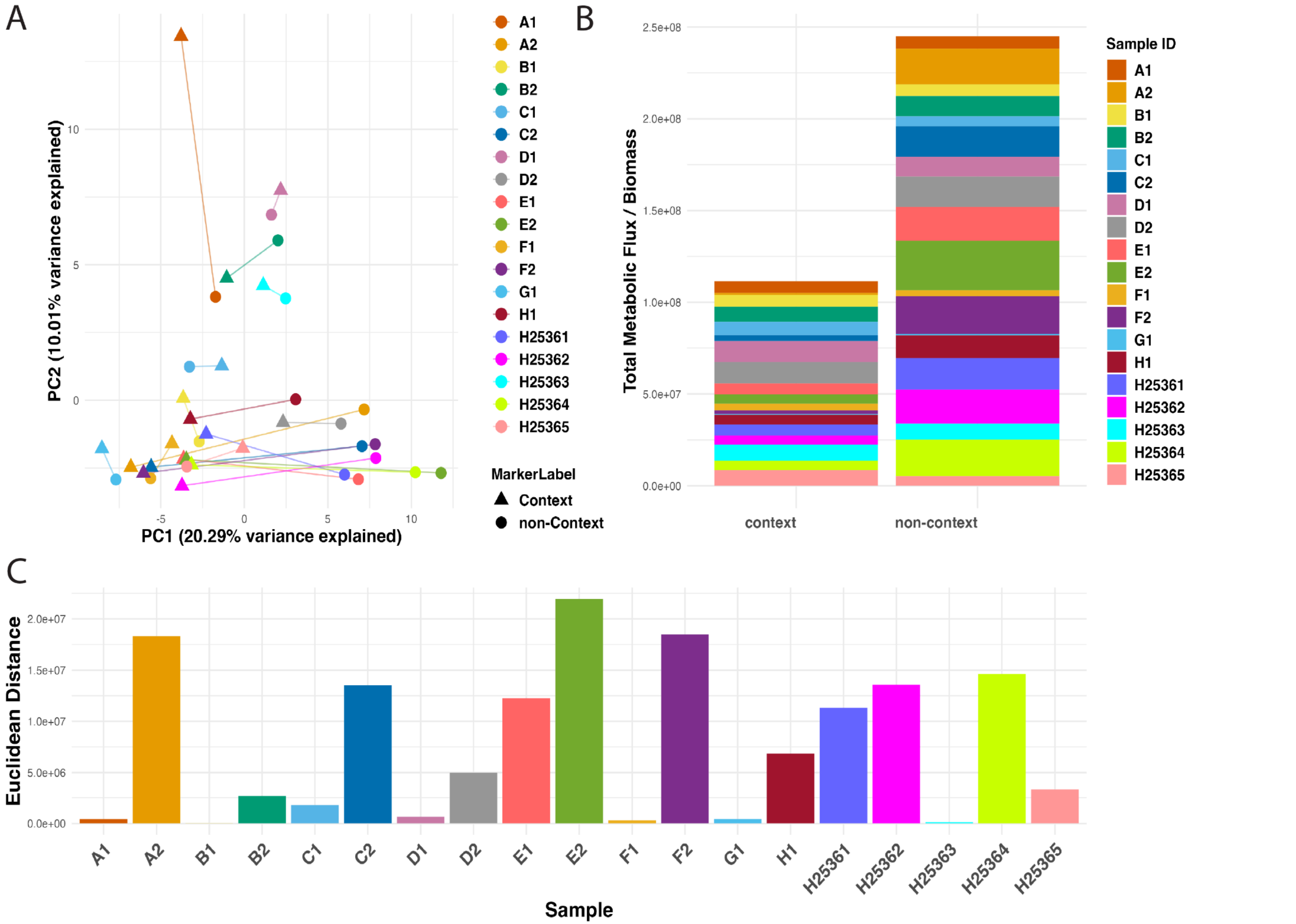
A
B
C
PC2 (10.01% variance explained)
PC1 (20.29% variance explained)
10
5
0
-5
0
5
10
A1
A2
B1
B2
C1
C2
D1
D2
E1
E2
F1
F2
G1
H1
H25361
H25362
H25363
H25364
H25365
MarkerLabel
Context
non-Context
Total Metabolic Flux / Biomass
2.5e+08
2.0e+08
1.5e+08
1.0e+08
5.0e+07
0.0e+00
context
non-context
Sample ID
Euclidean Distance
2.0e+07
1.5e+07
1.0e+07
5.0e+06
0.0e+00
Sample

**Supplementary Figure S3: Comparative analysis of metabolic flux across non-context-specific and context-specific microbiome models**

*By describing differences in metabolic flux normalized by community biomass between non-context-specific versus context-specific community modeling, changes in metabolites due to methodology can be observed. (A) A Principal Component Analysis (PCA) plot illustrates metabolic flux distribution across different patient samples and modeling methodologies. (B)* Total metabolic flux between context and non-context-specific models. (C) Euclidean distances between non-context-specific and context-specific conditions for each sample, indicating the differential impact of context on metabolic flux or biomass. Differences in height represent the disparity in metabolic activity, with larger distances reflecting significant methodological effects on the microbial community's metabolic outcomes.
